# Reduced placental transfer of antibodies against microbial and vaccine antigens in HIV-infected women in Mozambique

**DOI:** 10.1101/2020.08.05.237503

**Authors:** Selena Alonso, Marta Vidal, Gemma Ruiz-Olalla, Raquel González, M. Nelia Manaca, Chenjerai Jairoce, Miquel Vázquez-Santiago, Reyes Balcells, Anifa Vala, María Ruperez, Pau Cisteró, Laura Fuente-Soro, Marta Cova, Evelina Angov, Arsenio Nhacolo, Esperança Sevene, John J. Aponte, Eusébio Macete, Ruth Aguilar, Alfredo Mayor, Clara Menéndez, Carlota Dobaño, Gemma Moncunill

## Abstract

Antibody transplacental transfer is essential for conferring protection in newborns against infectious diseases. This transfer may be affected by gestational age and maternal infections, although the effects are not consistent across studies. We measured total IgG and IgG subclasses by quantitative suspension array technology against fourteen pathogens and vaccine antigens, including target of maternal immunization, in 341 delivering HIV− and HIV+ mother-infant pairs from southern Mozambique. Maternal antibody levels were the main determinant of cord antibody levels. HIV broadly reduced the placental transfer and cord levels of IgG and IgG1, but also IgG2 to half of the antigens. *Plasmodium falciparum* exposure and prematurity were negatively associated with cord antibody levels and placental transfer but this was antigen-subclass dependent. These findings suggest maternal infections may impact the efficacy of maternal immunization and confirm the lower transfer of antibodies as one of the causes underlying increased susceptibility to infections in HIV-exposed infants.

## Introduction

Each year, 2.6 million deaths occur during the neonatal period, being infectious diseases the leading cause of mortality, particularly in low-income countries [1, 2]. Newborns are highly vulnerable to pathogens due to their functional immunological differences from adults as a result of living in a semi-allogeneic sterile environment, where exposure to microbial antigens is limited [3–6]. For example, microorganisms such as respiratory syncytial virus (RSV) are generally asymptomatic or cause mild disease in adults but induce acute bronchiolitis, viral pneumoniae and croup in infants, being those between 2 and 6 months of age at the highest risk, especially in low-income countries [7, 8].

Newborns mostly rely on the protection elicited by maternal antibodies transferred across the placenta, which provide passive immunity against common pathogens [9]. Neonatal and child immunization is essential for conferring protection in newborns and infants against vaccine-preventable diseases [10–12]. Vaccination is among the most cost-effective public health measures worldwide [13], and regions with high rates of infant morbidity and mortality like sub-Saharan Africa benefit from the implementation of the Expanded Program of Immunization (EPI) [14]. Nevertheless, acquisition of immunity from vaccination is not immediate and vaccines are not available for all infectious diseases. At present, only three vaccines are being administered at birth in some countries: Bacillus Calmette-Guérin (BCG), hepatitis B virus (HBV) and oral polio vaccine (OPV) [10, 15].

Transplacental transfer of antibodies occurs in utero and it is facilitated by neonatal fragment crystallisable (Fc) region receptor (FcRn), expressed in the human syncytiotrophoblast [16]. Only IgG is transferred across the placenta, being foetal IgG concentrations higher at the third trimester [17], although some studies suggest that maternal IgE is also transferred to the foetus as IgG/IgE complexes [18]. IgG subclasses have different affinities for the FcRn receptor leading to differences in the efficiency of transfer [19]; the greatest transport occurs for IgG1, followed by IgG4, IgG3, and finally IgG2 [20].

To be effective, the transferred IgG must reach protective levels after birth. Maternal immunization is a good strategy to prevent newborn infections, ensuring a sufficient transfer of protective antibodies to the neonate [21]. Maternal vaccination against tetanus, pertussis and influenza has been implemented in many populations and has been effective protecting young infants from these pathogens [22–24], and could be used to protect newborns from RSV [25].

A number of factors have been associated with IgG placental transfer and cord levels, such as maternal antibody concentrations, gestational age, placental integrity, maternal infections and the antigen specificity [26–30], but inconsistently. Placental malaria (PM) has been shown to reduce transplacental transfer of antibodies against tetanus, measles, *Streptococcus pneumoniae* (*S. pneumoniae*), herpes simplex virus type 1 (HSV-1), RSV and varicella-zoster virus (VZV) [26, 31–33]. However, other studies have shown no impact of PM on transplacental transfer of tetanus, *S. pneumoniae, Haemophilus influenzae type b* (*Hib*), diphtheria, measles or RSV antibodies [31, 33–36]. The effect of maternal HIV infection is also controversial. Some studies demonstrated that HIV infection leads to a reduction of the transplacental transfer of *Hib*, pertussis, pneumococcus, measles, tetanus and *Plasmodium falciparum* (*P. falciparum*) specific antibodies [26, 27, 37–39], but others have shown no effect [31, 36, 39–41]. Therefore, there are probably confounding variables that should be considered.

In our study, we wanted to assess the impact of different factors, including maternal HIV infection and malaria in pregnancy on the placental transfer and cord levels of IgG and IgG subclasses to a broad range of highly prevalent microbial and vaccine antigens in a sub-Saharan African country, including: *Corynebacterium diphtheriae* (*C. diphtheriae*), *Clostridium tetani* (*C. tetani*)*, Bordetella pertussis* (*B. pertussis*)*, Hib, S. pneumoniae, Shigella dysenteriae* (*S. dystenteriae*)*, Vibrio cholerae* (*V. cholerae*), hepatitis B virus (HBV), measles, RSV, *Cryptosporidium parvum* (*C. parvum*)*, Giardia intestinalis* (*G. intestinalis*) and *P. falciparum*. A better understanding of factors affecting cord IgG levels will help designing better preventive measures and strategies for maternal and child health.

## Results

### Description of participants

A total of 341 women (197 HIV-uninfected and 144 HIV-infected) participated in the study (Table 1). HIV-infected women were older than the HIV-uninfected and there were more primigravidae among the HIV-uninfected. HIV-infected women had significantly more anaemia than the HIV-uninfected. There were no significant differences in birth weight or prematurity between infants born to HIV-infected and those born to HIV-uninfected women. Among the 155 infants born from HIV-infected women, 8 tested HIV-positive at 6 weeks of age by polymerase chain reaction (PCR) analysis performed following national guidelines. Placental histology was performed on 307 samples from study participants, of which 3 had acute PM and 8 past PM. In total, 20 women had PM (positive at placental blood, by microscopy or PCR at delivery, or acute or past PM by histology), but there were no differences by HIV infection. Peripheral malaria (positive at peripheral blood by microscopy or PCR at any of the visits during pregnancy) was detected in 51 women, but there were no differences by HIV infection. Finally, *P. falciparum* exposure was lower among HIV-infected women.

**Table 1:** Characteristics of study participants.

|  | All<br>N=341 | HIV-uninfected<br>N=197 | HIV-infected<br>N=144 | p-value<sup>a</sup> |
| --- | --- | --- | --- | --- |
| Age<sup>a</sup> (years median [IQR]) | 25.0 [19.0; 29.0] | 21.0 [18.0; 28.0] | 27.0 [22.0; 31.0] | < 0.001 |
| Gravidity (n, %) |  |  |  | < 0.001 |
| *Multigravidae* | 259 (76.0) | 128 (65.0) | 131 (91.0) |  |
| *Primigravidae* | 82 (24.0) | 69 (35.0) | 13 (9.0) |  |
| Maternal haemoglobin (n, %) |  |  |  | 0.025 |
| Anaemia (< 11 g/dL) | 208 (61.5) | 109 (56.2) | 99 (68.8) |  |
| Normal (≥ 11 g/dL) | 130 (38.5) | 85 (43.8) | 45 (31.2) |  |
| Birth weight (n, %) |  |  |  | NS |
| Low (< 2500 g) | 29 (8.5) | 17 (8.6) | 12 (8.33) |  |
| Normal (≥ 2500 g) | 312 (91.5) | 180 (91.4) | 132 (91.7) |  |
| Prematurity (n, %) |  |  |  | NS |
| No (≥ 37 weeks) | 312 (94.3) | 181 (95.3) | 131 (92.9) |  |
| Yes (< 37 weeks) | 19 (5.7) | 9 (4.7) | 10 (7.1) |  |
| Treatment |  |  |  | < 0.001 |
| MQ | 71 (20.9) | 0 (0.0) | 71 (49.7) |  |
| MQ full | 68 (20.8) | 68 (34.5) | 0 (0.0) |  |
| MQ split | 73 (21.5) | 73 (37.1) | 0 (0.0) |  |
| Placebo | 72 (21.2) | 0 (0.0) | 72 (50.3) |  |
| SP | 56 (16.5) | 56 (28.4) | 0 (0.0) |  |
| ART at baseline (n, %) |  |  |  | NP |
| No | 24 (7.1) | – | 24 (17.1) |  |
| Yes | 116 (34.4) | – | 116 (82.9) |  |
| CD4+ T cell counts (n, %) |  |  |  | NP |
| Lower (< 350 c/μL) | 40 (12.3) | – | 40 (31.2) |  |
| Higher (≥ 350 c/μL) | 88 (27.1) | – | 88 (68.8) |  |
| HIV viral load (copies/mL) |  |  |  | NP |
| < 400 | 21 (6.4) | – | 21 (16.0) |  |
| (400–999) | 41 (12.5) | – | 41 (31.3) |  |
| (1000–9999) | 48 (14.6) | – | 48 (36.6) |  |
| > 9999 | 21 (6.4) | – | 21 (16.0) |  |
| Placental malaria<sup>b</sup> (n, %) |  |  |  | NS |
| No | 321 (94.1) | 184 (93.4) | 137 (95.1) |  |
| Yes | 20 (5.9) | 13 (6.6) | 7 (4.9) |  |
| Peripheral malaria<sup>c</sup> (n, %) |  |  |  | NS |
| No | 290 (85.0) | 165 (83.8) | 125 (86.8) |  |
| Yes | 51 (15.0) | 32 (16.2) | 19 (13.2) |  |
| *P. falciparum* exposure (log<sub>10</sub> MFI IgG) | 5.27 [5.19; 5.34] | 5.29 [5.21; 5.35] | 5.26 [5.18; 5.33] | 0.011 |

For numerical variables, the median and first and third quantile, in brackets, are given. For the categorical variables the number of individuals for each group and percentages, in parentheses, are given.
<sup>a</sup> For the age, the Wilcoxon-Mann-Whitney test was used to compare differences between median values. For the categorical variables, the Chi-square test was used.
<sup>b</sup> Placental malaria was considered positive if there was any evidence of *P. falciparum* placental parasitaemia by any method.
<sup>c</sup> Peripheral malaria was considered positive if there was any evidence of *P. falciparum* peripheral parasitaemia by any method.
The statistical significance was considered when *p*-value < 0.05; MQ, mefloquine; NS, not significant; NP, not-performed tests; SP, sulfadoxine-pyrimethamine.

### Profile of antibody levels in cord blood

We first performed principal component analysis (PCA) of the cord antibody levels and maternal antibody levels separately, including IgG, IgG1, IgG2, IgG3 and IgG4 to all antigens tested, to reduce the dimensionality of the data and get insights into the overall antibody patterns. Cord and maternal PCA looked very similar (data not shown). Cord antibody responses clearly clustered by IgG subclasses (Fig. 1a) and antigens (Fig. 1b) suggesting different antibody profiles depending on the antigen specificity. IgG and IgG1 clusters were closer, showing similar responses, whereas IgG4 and IgG3 were the most distant. *Hib* cluster was clearly separated from the rest indicating a different antibody profile. Consistently, median IgG and IgG1 levels were higher than the rest of IgG subclasses and were both similar between them for most of the antigens with the exception of *Hib* (Fig. 1c). IgG2 had lower median levels than IgG1, followed by IgG3 and the lowest levels were shown for IgG4.

**Fig. 1:**
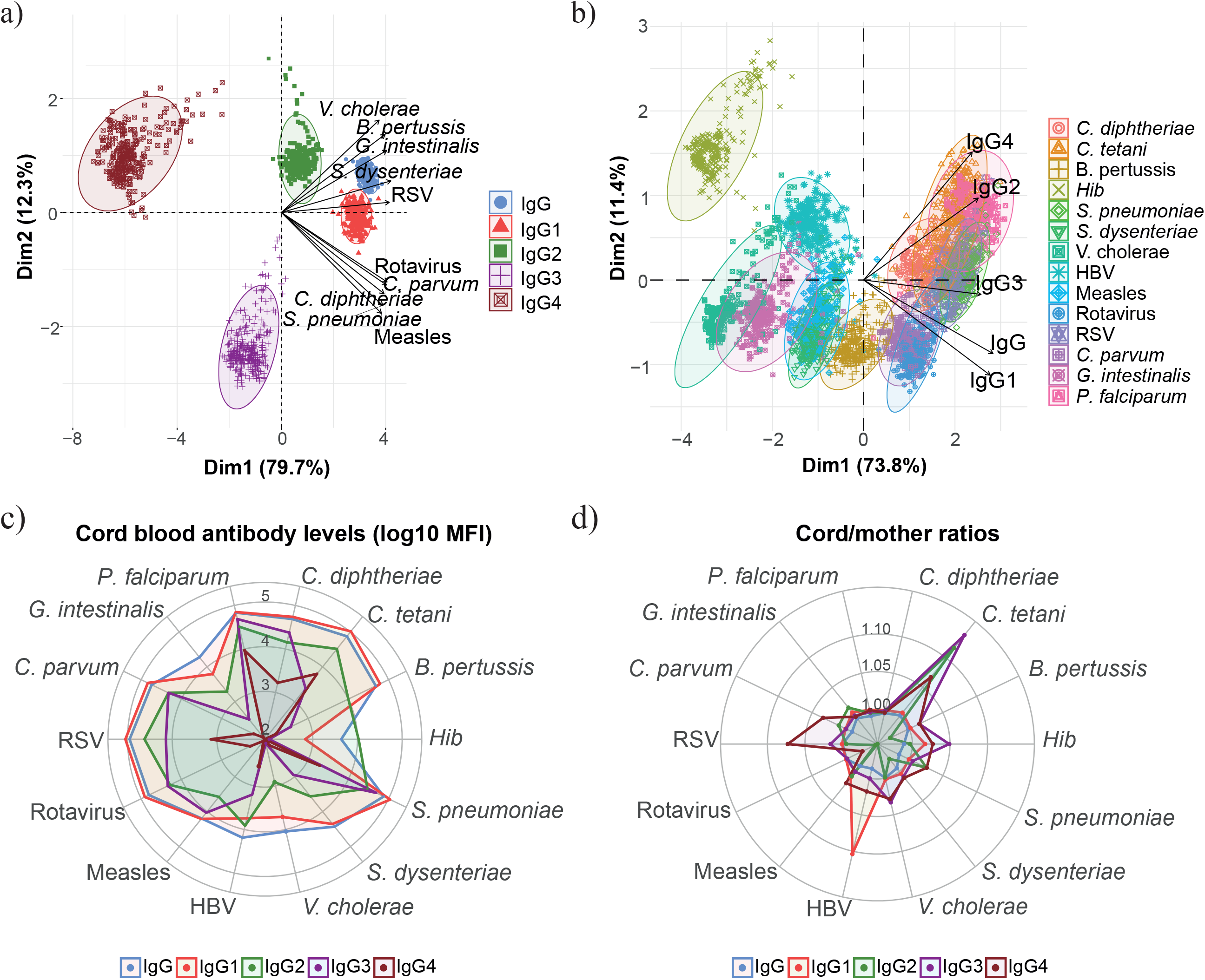
Overview of levels of IgG and IgG subclasses to all pathogen antigens. a) Principal component analysis (PCA) plots of IgG and IgG subclass levels against all antigens clustered by subclass type. b) PCA plots of IgG and IgG subclass levels clustered by antigen type. The two principal components (Dim 1, Dim2) that explained the highest percentage of the variance of the data (percentage in parenthesis) were chosen for representation. Representation of the a) 10 b) 5 first variables that contributed to the principal components c) Medians of IgG and IgG subclass levels (log_10_ MFI) in cord blood for each antigen d) Median IgG and IgG subclass placental transfer for each antigen, represented as the cord/mother ratios. Source files of the medians of each antigen/subclass are available in the Figure 1-source data 1.

We also determined the placental transfer of antibodies, measured as the ratio of cord blood levels to the maternal levels. The transfer efficiency was greatest for IgG1, IgG3 or IgG4 depending on the antigen: IgG1 for *C. diphtheriae, P. falciparum,* HBV and rotavirus, IgG3 for *B. pertussis, C. tetani, Hib* and *V. cholerae* and IgG4 for *C. parvum, S. dysenteriae,* measles and RSV. The less efficiently transferred subclass was IgG2 for most of the antigens, with the exception of *G. intestinalis* and *S. pneumoniae* for which IgG2 was the greatest (Fig. 1d).

### Altered maternal and cord blood antibody levels in HIV-infected women

HIV-infected and HIV-uninfected women did not show significant differences between antibody levels except for *C. tetani* (IgG and IgG1), *S. pneunomiae* and RSV (IgG2), and *C. diphtheriae* and *P. falciparum* (IgG4), with lower antibody levels in HIV-infected compared to HIV-uninfected women (Fig. 2a-2d and Figure 2-figure supplement 1). In contrast, higher *G. intestinalis* and HBV IgG levels were found in HIV-infected women (Fig. 2a).

**Fig. 2:**
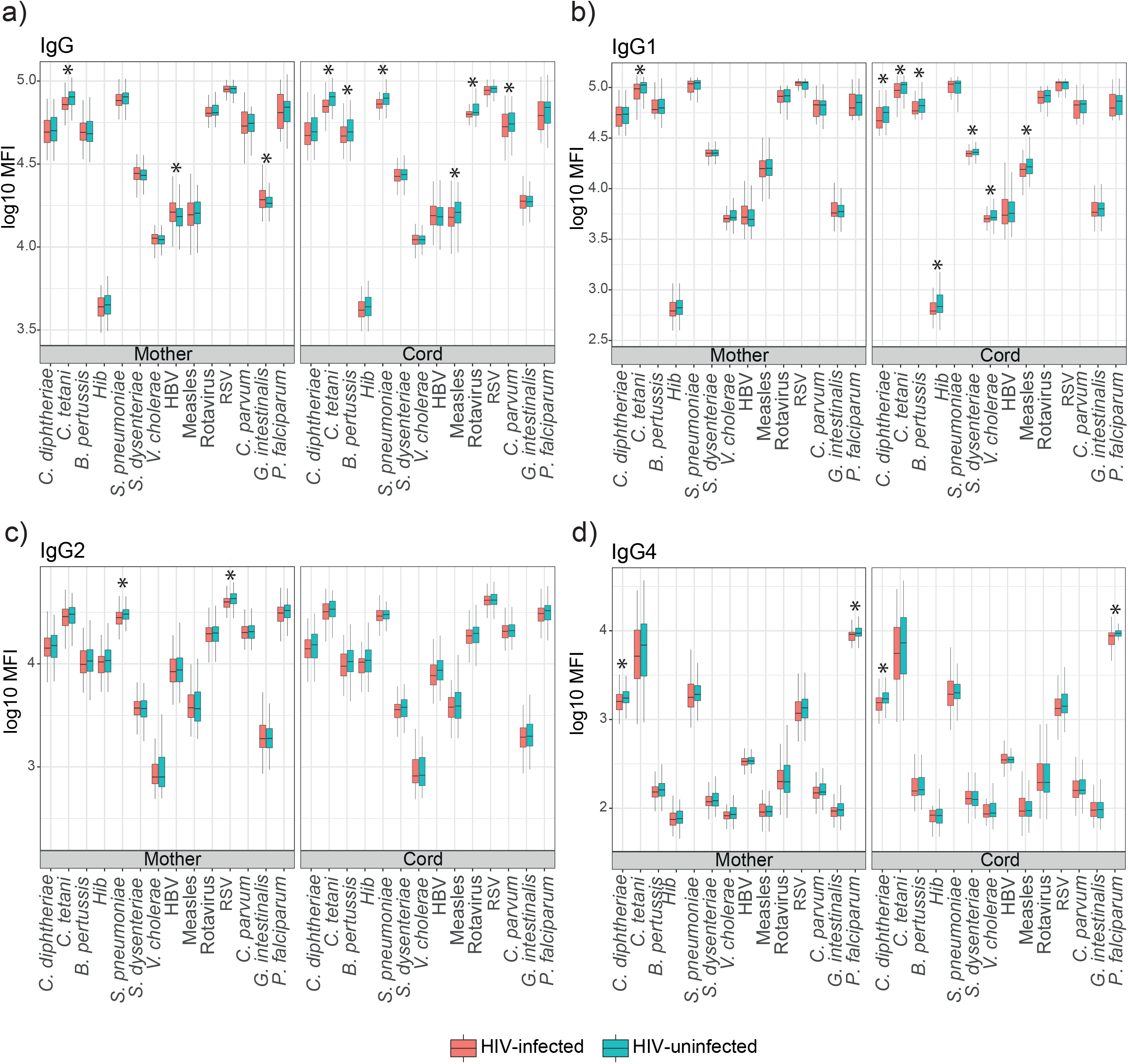
Mother and cord blood antibody levels (log_10_ MFI) in HIV-infected and HIV-uninfected women. Boxplots illustrate the medians and the interquartile range for IgG (a) and IgG1 (b), IgG2 (c) and IgG4 (d) subclasses. Levels between HIV-infected and HIV-uninfected women were compared by parametric Wilcoxon-Mann–Whitney test and p-values were adjusted for multiple testing by the Benjamini-Hochberg approach. Statistically significant differences between HIV infected and uninfected women levels are highlighted with an asterisk. Red represents HIV-infected women and blue HIV-uninfected women. Source files of the mother and cord levels of each antigen/subclass are available in the Figure 2-source data 1.

IgG cord blood levels were lower in HIV-infected than HIV-uninfected women for *C. tetani, B. pertussis, S. pneumoniae,* measles, rotavirus and *C. parvum*. Similarly, HIV-infected women had lower IgG1 cord levels for *C. diphtheriae, C. tetani, B. pertussis, Hib, S. dysenteriae, V. cholerae* and measles for IgG1 (Fig. 2b). Lower *C. diphtheriae and P. falciparum* IgG4 levels were also found in cord blood of HIV-infected than HIV-uninfected women, whereas no differences were observed between groups for IgG2 and IgG3 (Fig 2c-2d and Figure 2-figure supplement 1).

### Altered placental transfer of antibodies in HIV-infected women

Placental transfer of IgG and IgG1 was significantly lower in HIV-infected women for all antigens except for *Hib* and *V. cholerae* (IgG) and *C. tetani, S. pneumoniae, V. cholerae* and RSV (IgG1) (Fig. 3a-3b, Figure 3-figure supplement 1 and Figure 3-figure supplement 2). For IgG2, only *G. intestinalis, B. pertussis* and HBV had significantly lower transfer in HIV-infected women, while *S. pneumoniae* and RSV had higher transfer in HIV-infected women (Fig. 3c and Figure 2-figure supplement 3). For IgG3, only *C. tetani* antibodies had a significantly lower transfer in HIV-infected compared to HIV-uninfected women (Fig. 3d and Figure 3-figure supplement 4). No significant differences in placental transfer between the two groups were found for IgG4 (Figure 3-figure supplement 5).

**Fig. 3:**
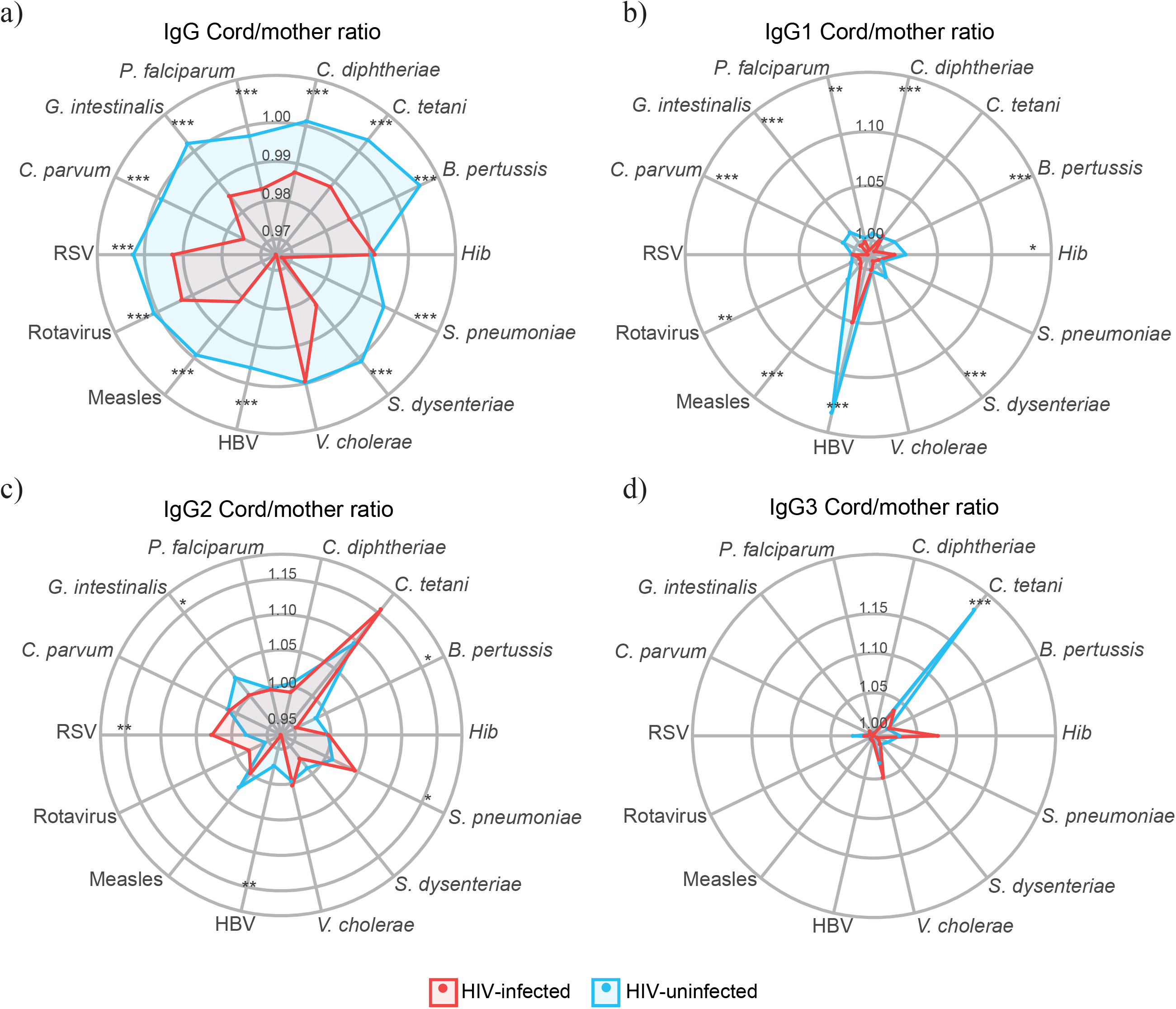
Antibody placental transfer in HIV-infected and HIV-uninfected women. Radar charts representing the medians of each analyte antibody cord/mother ratio in HIV-infected and HIV-uninfected women for IgG (a) and IgG subclasses (b-d). Ratios between HIV-infected and uninfected women were compared by parametric Wilcoxon-Mann-Whitney test and p-values were adjusted for multiple testing by the Benjamini-Hochberg approach. Statistically significant differences between HIV-infected and uninfected women ratios are highlighted with an asterisk. *** = p-val < 0.0001, ** = p-val < 0.001, * = p-val < 0.01. Red represents HIV-infected women and blue HIV-uninfected women. Source files of the medians and p-values of each antigen/subclass are available in the Figure 3-source data 1.

### Factors associated with cord blood levels of IgG and IgG subclasses

Maternal antibodies, HIV infection and *P. falciparum* exposure were the only variables with a clear general impact on univariable models and were selected for multivariable models, in which maternal antibody levels had the strongest positive correlation with cord antibody levels for all the antigens and subclasses (Fig. 4a). However, the effect of maternal antibody levels was more variable for IgG3-4 than for IgG and IgG1 subclasses. On average, a 10% increase in maternal IgG levels was associated with 8.1% to 9.7% increases in IgG cord blood levels, depending on the antigen. For IgG subclasses, a 10% increase in maternal antibody levels was associated with increases of cord blood levels from 7.6% to 10.9% for IgG1, 5.4% to 9.6% for IgG2, 5% to 9.9% for IgG3 and 5.3% to 9.3% for IgG4.

**Fig. 4:**
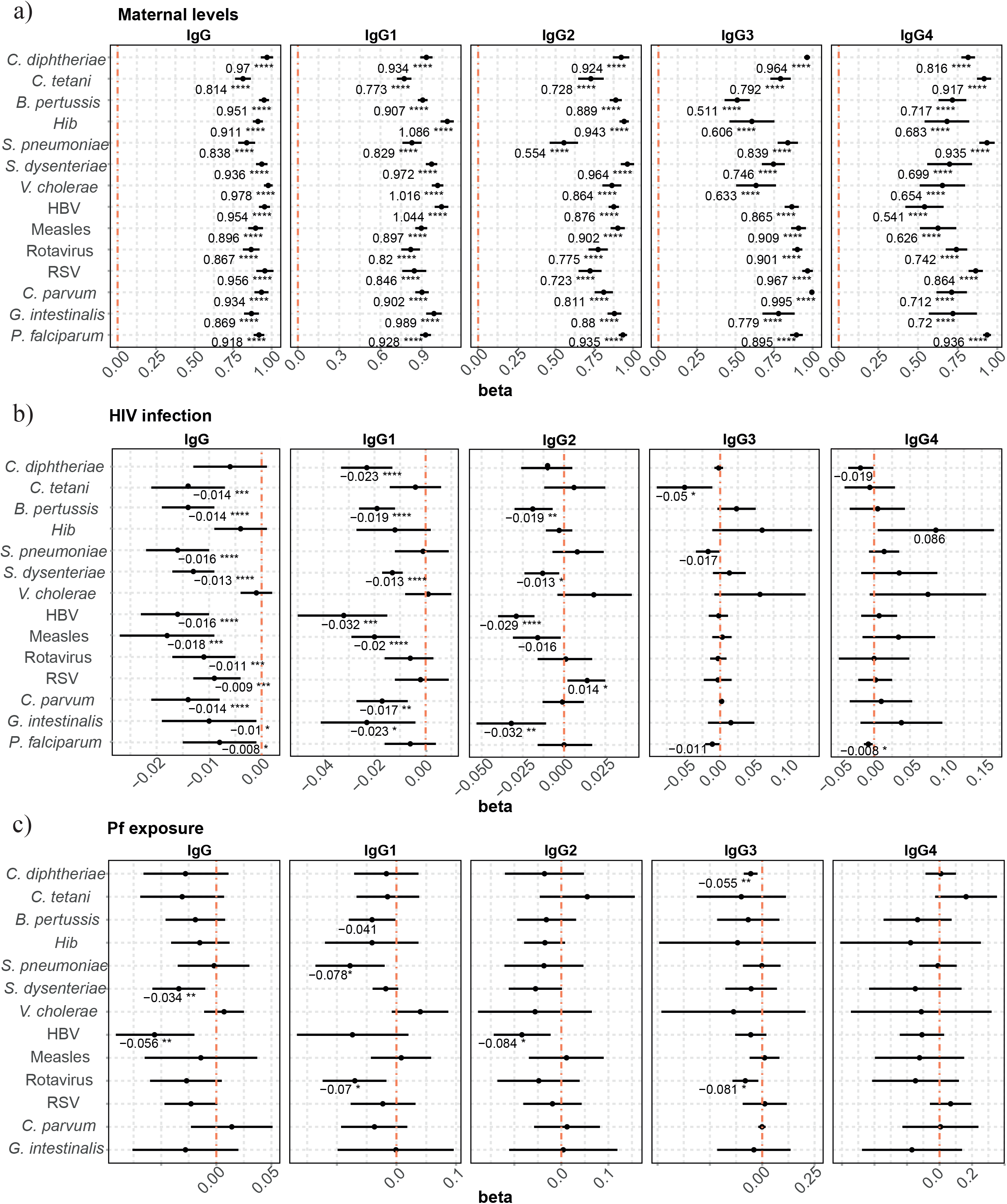
Factors associated with IgG and IgG subclass levels in cord blood. Forest plots show the effect of a) maternal antibody levels, b) HIV infection and c) *P. falciparum* exposure (Pf exposure) on cord blood levels of IgG and IgG subclasses, for all the antigens tested. Beta values, representing the increase or decrease of cord blood levels (log_10_MFI) were obtained from multivariable regression models using cord blood (log_10_MFI) levels as the outcome. Beta values are shown when raw p-vals are significant. Asterisks are shown when adjusted p-vals by Benjamini-Hochbert are significant **** = p-val < 0.0001, *** = p-val < 0.001, ** = p-val < 0.01, * = p-val < 0.05. Source files of the multivariable model are available in the Figure 4-source data 1.

Maternal HIV infection (Fig. 4b) had a negative effect on IgG cord blood levels to all antigens, except for *C. diphtheriae, Hib* and *V. cholerae*. HIV infection was associated with a 2.1% to 4.1% reduction in the IgG cord blood levels. For IgG1, HIV infection negatively impacted cord blood levels against *C. diphtheriae, B. pertussis, S. dysenteriae*, HBV, measles, *C. parvum* and *G. intestinalis* (2.9% to 7.1% reduction). For IgG2, an HIV negative effect was observed against *B. pertussis, S. dysenteriae*, HBV and *G. intestinalis* (2.9% to 7.1% reduction), whereas HIV was associated with an increase of 3.3% of IgG2 to RSV. Finally, we only detected a negative effect of HIV infection on IgG3 levels to *C. tetani* (1.1% reduction) and IgG4 to *P. falciparum* (1.8% reduction).

*P. falciparum* exposure was negatively associated with cord blood IgG levels against *S. dysenteriae* and HBV, IgG1 against *S. pneumoniae* and rotavirus, IgG2 against HBV and IgG3 against *C. diphtheriae* and rotavirus (Fig. 4c). Depending on the IgG sublcass and antigen, 10% increseas in *P. falciparum* exposure reduced the cord blood levels from 0.3% to 0.8%.

Previous studies suggest that PM rather than peripheral malaria affect transplacental transfer of antibodies and lead to adverse outcomes due to the damaged placental tissue [35, 42, 43]. Therefore, we explored the effect of PM on cord blood levels and placental transfer instead of *P. falciparum* exposure despite the low number of women with any evidence of PM. When analysing HIV-infected women only, PM was associated with lower *B. pertussis* IgG1, *C. diphtheriae* IgG2 and HBV IgG3 levels in cord blood (Figure 4-figure supplement 1).

Prematurity (Fig. 5a), previously shown to have a detrimental effect on placental transfer of antibodies [44], increased the quality (AIC) of some of the above multivariable models. Prematurity was associated with lower cord blood IgG levels against *Hib* (4.2% reduction compared with on-term cord blood levels)*, V. cholerae* (2.3% reduction), measles (5.8% reduction) and *C. parvum* (3,8% reduction without statistical significance after adjusting for multiple testing)

**Fig. 5:**
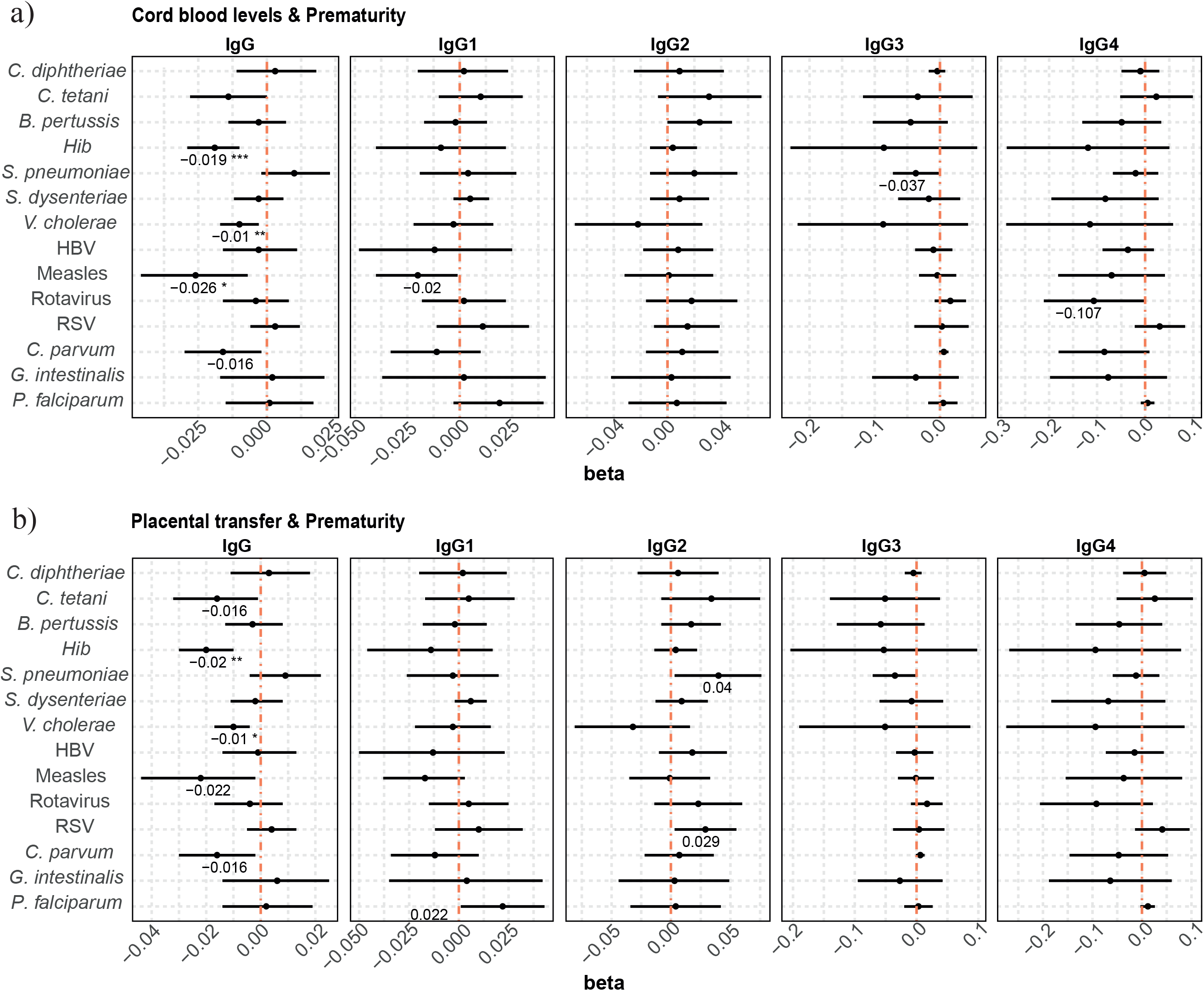
Association of prematurity with cord blood levels and placental transfer of IgG and IgG subclasses. Forest plots show the effect of a) prematurity and cord blood levels and b) prematurity and transplacental transfer of IgG and IgG subclasses, for all the antigens tested. Cord antibody levels are represented in log_10_MFI. Placental transfer is represented as cord/mother ratio (log_10_). Beta values are shown when raw p-vals are significant. Asterisks are shown when adjusted p-vals by Benjamini-Hochbert are significant. **** = p-val < 0.0001, *** = p-val < 0.001, ** = p-val < 0.01, * = p-val < 0.05. Source files of the multivariable model are available in the Figure 5-source data 1.

The rest of the variables (age, maternal anaemia, gravidity, low birth weight, IPTp treatment, seasonality; and CD4^+^ T cell counts, ART and viral load for HIV-infected women) did not provide an added value to the multivariable models. Univariable models did not show a consistent effect of any variable across antigens or IgG subclasses, but some significant associations were found for age and gravidity (Supplementary Material 1).

### Factors associated with placental transfer of IgG and IgG subclasses

In multivariable models including HIV infection and *P. falciparum* exposure, HIV infection (Fig. 6a) was associated with a generalized reduced placental transfer of IgG and IgG1 (from 2.1% to 6.7% reduction depending on the antigen). HIV infection was also associated with a reduced transfer of IgG2 against *B. pertussis*, HBV and *G. intestinalis*, but was associated with an increase in IgG2 RSV transfer (5.4% increase). Although adjusted p-values were not significant, a similar trend of positive correlation was found for *S. pneumoniae* IgG2 and *Hib* and *V. cholerae* IgG3 and IgG4.

**Fig. 6:**
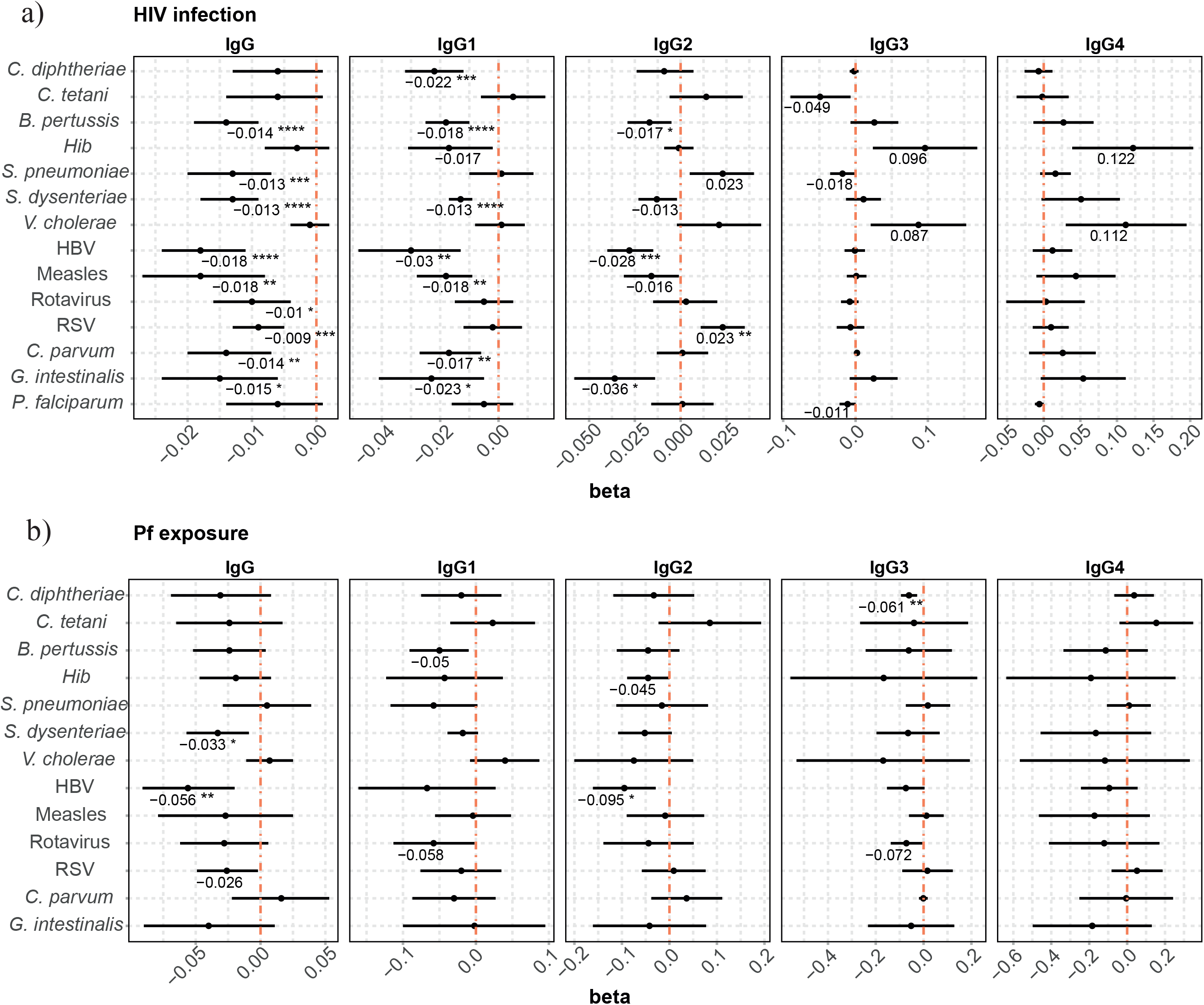
Factors associated with IgG and IgG subclass placental transfer. Forest plots show the effect of a) HIV infection and b) *P. falciparum* exposure (Pf exposure) on placental transfer of IgG and IgG subclasses, for all the antigens tested. Beta values are shown when raw p-vals are significant. Asterisks are shown when adjusted p-vals by Benjamini-Hochbert are significant. **** = p-val < 0.0001, *** = p-val < 0.001, ** = p-val < 0.01, * = p-val < 0.05. Source files of the multivariable model are available in the Figure 6-source data 1.

*P. falciparum* exposure (Fig. 6b) had a negative effect on the placental transfer of antibodies for some antigens. An increase of 10% in *P. falciparum* exposure reduced the placental transfer of IgG against *S. dysenteriae* and HBV by 0.3% and 0.5%, respectively, IgG2 against HBV by 0.9% and IgG3 against *C. diphtheriae* by 0.6%.

PM, in contrast to *P. falciparum* exposure, did not have any impact on transplacental transfer of antibodies in exploratory analyses and did not improve any of the models, although it had a similar trend of correlation on IgG. Nevertheless, PM was associated with a diminished placental transfer on IgG1 *B. pertussis* among the HIV-positive subset of women (Figure 6-figure supplement 1).

When prematurity was added to the multivariable models, this additional covariable had a negative effect on placental transfer of *Hib* and *V. cholerae* IgG antibodies (4.5% and 2.3% reduction in premature vs on-term newborns, respectively) (Fig. 5b).

The rest of variables were not added to the placental transfer multivariable models, because almost none of the models improved when included. Similar to cord blood levels ones, the placental transfer univariable models did not show a consistent effect of any variable not included in the multivariable models across antigens or IgG subclasses (Supplementary Material 1).

## Discussion

Our comprehensive analysis of maternal and cord plasma IgG and IgG subclasses against a wide range of microbial and vaccine antigens allowed a depth immunoprofiling, that is essential to decipher the mechanisms affecting antibody placental transfer and maternal and newborn immunity in women chronically exposed to pathogens. We confirmed that the main determinant of cord IgG and IgG subclass levels are the maternal corresponding antibody levels, and that maternal HIV infection is associated with a generalized diminished IgG levels in the cord due to low maternal levels but also to a broadly reduction of IgG and IgG1 placental transfer.

Maternal and cord blood antibody levels are usually correlated in many studies, suggesting that maternal levels are the main determinant for transfer efficiency [9, 45, 46]. However, the effect of HIV infection on placental transfer has not been consistently assessed and the few studies looking at its effect on maternal and cord blood levels mainly focussed on total IgG. Our results showed that HIV infection reduced the IgG maternal levels for some antigens, the cord blood levels overall, and also had a negative effect on transplacental transfer of IgG antibodies. It is interesting that although we found higher maternal HBV and *G. intestinalis* IgG levels among HIV-infected women, cord blood levels and transplacental transfer were lower than in HIV-uninfected women. Higher maternal antibody levels against these pathogens in HIV-infected women may be due to an increased susceptibility to co-infections with these pathogens, as described before [47–49], but it seems that they are not being transferred as efficiently as in HIV-uninfected women. This could be due to hypergammaglobulinemia, demonstrated to be common among HIV-infected individuals [50] and previously shown to impair transplacental transfer of antibodies [9, 33].

Our results are consistent with previous studies reporting that HIV infection led to a reduction of the cord blood levels and transplacental transfer of total IgG against *B. pertussis* [40, 41]*, C. tetani* [26, 38, 40, 41]*, S. pneumoniae* [31, 38, 41, 51], RSV [52, 53] and measles [37, 38]. Some studies also found a negative effect on *Hib* [27, 40, 51] that is not appreciated in our study (although we found reduced IgG1 levels in cord in univariable analyses). However, our results differ from other studies that did not find any effect of HIV status on IgG levels against *C. diphtheriae* [36]*, C. tetani* [31, 36], *S. pneumoniae* [53], HBV [36] and measles [31, 36].

IgG subclasses may be differently elicited depending on the pathogen, the antigen or the epitope [54] and the efficiency of the antibody placental transfer is different for each subclass due to differential affinity of the receptors FcRn. Furthermore, the Fc region of IgG, that mediates effector functions, vary between IgG subclasses, conferring them different roles during infection and pathogen clearance [55]. We found that HIV infection reduced mainly IgG1 cord levels due to an HIV impairment of the transplacental transfer, similarly to IgG. Interestingly, maternal HIV infection increased the placental transfer of IgG2 to *S. pneumoniae* and RSV, although in multivariable models it was only significant for RSV. We also found that HIV infection had a positive effect on RSV IgG2 cord blood levels, although IgG2 maternal levels were lower among HIV-infected women. To our knowledge an increased placental transfer by HIV infection has not been described before. This may have implications for maternal immunization with RSV vaccines under development.

The efficacy of IgG placental transfer also depended on the antigen. IgG1, IgG3 or IgG4 transferred better than IgG2, except for *S. pneumoniae* and *G. intestinalis*, for which IgG2 transfer was higher. This was unexpected because it has been previously described that the greatest transport occurs for IgG1, followed by IgG4, IgG3, and finally IgG2 [9, 20]. However, IgG1 levels were the highest for almost all antigens in cord blood, probably because the overall higher levels of this IgG subclass in maternal blood. One exception was *Hib* that presented higher IgG2 cord levels than IgG1, although IgG1 transplacental transfer was higher than IgG2 consistently with previous studies [56]. The mothers had a IgG2-predominant response to *Hib*, and consequently higher IgG2 than IgG1 levels were found in cord blood as previously described [57, 58].

Regarding other variables, our results did not show any significant association between CD4^+^ T cell counts or HIV viral load on cord blood levels and transplacental transfer of antibodies. Even though these results agree with previous studies that did not find any associations [40, 51, 59], other studies described that lower CD4^+^ T cell counts and higher HIV viral load led to a reduction on the transfer of some pathogen-specific antibodies and vaccines such as measles and *S. pneumoniae* [35, 60, 61]. Some studies described that HIV-infected women receiving ART transferred higher pathogen-specific antibodies than those who were not under ART [59] or who initiated it during pregnancy [62]. However, in our cohort we did not find any significant associations in regards to ART.

At the time of the study, malaria transmission intensity was very low in the area and only a few women had active malaria during pregnancy. Nonetheless, we found a negative correlation between *P. falciparum* exposure and both placental transfer and cord blood antibody levels for some antigens and IgG subclasses. Previous studies are contradictory, as some found that PM led to a reduction of the transplacental transfer of some pathogen-specific IgG to *C. tetani* [32], measles [33, 37], RSV [35] and *S. pneumoniae* [31], but others did not find any effect for IgG against *C. diphtheriae* [35, 36], *C. tetani* [26, 33, 36]*, Hib* [35], HBV [36], measles [36] RSV [34] and *S. pneumoniae* [35]. Discrepancies between studies could be due to the different study areas, with different prevalence of malaria and study sample sizes, different type of antigens used in the studies, the different sensibilities among the serological methods used, different exposure to the pathogens tested, and other co-infections.

We found prematurity to be associated with lower cord blood IgG levels and placental transfer for some antigens, as previous studies have shown [44, 63, 64], although the effect was not consistent among subclasses. It has already been reported that the greatest transport occur in the third trimester of gestation [17], and due to this fact, preterm infants may have lower amounts of transplacental IgG than term infants.

Our results are important for maternal immunization implementation in settings with a high prevalence of HIV infection. In our study cohort, the only vaccine given during pregnancy was tetanus. Although HIV infection was associated with lower maternal and cord blood tetanus toxoid IgG and IgG1 levels in univariable models, HIV did not affect cord blood IgG1 levels in multivariable models adjusted by maternal levels. Systemic tetanus vaccination during pregnancy has been implemented in Africa and has demonstrated a high efficacy [65]. Pertussis vaccination in pregnancy has also been implemented in some countries, but not in Africa. Acellular pertussis vaccine induces mainly IgG and IgG1 responses that are thought to confer protection [66, 67]. We found lower cord blood levels and a reduced placental transfer of IgG and IgG1 against *B. pertussis* among HIV-infected women and those exposed to *P. falciparum*. These results highlight the need for further studies assessing the impact of these infections on pertussis vaccine efficacy and antibody placental transfer when implemented in pregnant women from African countries. A current vaccine in development for maternal immunization is RSV [68]. Natural RSV infection seems to elicit an IgG1 and IgG2 response against the F protein, the major target of the host’s immune response [69] and of some vaccines in development [70]. Antibodies binding to the F protein were protective [71] and Palivizumab, an IgG1 monoclonal antibody against RSV F protein with neutralizing function, has shown to be effective [72]. Here, IgG and IgG1 against RSV F protein had the highest levels in cord blood compared to other subclasses, but HIV infection reduced IgG cord blood levels and placental transfer in multivariable models. Instead, IgG2 cord blood levels were increased by maternal HIV infection. Therefore, HIV infection could compromise the levels of RSV neutralizing antibodies transferred to the newborn and, consequently, diminish the effectivity of a RSV vaccine.

Unfortunately, we do not know what are the thresholds of antibody levels that confer protection in our study, therefore it is difficult to infer the clinical relevance of the reductions in antibody levels detected in cord blood from the-HIV infected women. A study in South Africa reported that the frequency of HIV-infected and HIV-uninfected pregnant women with protective antibody levels against pertussis, tetanus or HBV was similar, although the overall frequencies were low (32%, 41% and 30%, respectively) [40]. This same study demonstrated that the proportion of HIV-infected pregnant women reaching anti-*Hib* protective antibody levels was lower than HIV-uninfected women (35% vs 59%). Thus, for the implementation of maternal immunization programs, the effect of HIV infection and *P. falciparum* exposure must be taken into account, especially after demonstrating that both infections reduce the levels of antibodies in the cord blood and therefore may compromise vaccines protective effect in the newborn.

In conclusion, our results demonstrate that maternal HIV infection was associated with reduced levels of antibodies against a broad range of pathogens and vaccine antigens in cord blood. Part of this reduction in antibody levels was due to altered antibody levels in the mother, which are the main determinants of cord blood levels, but HIV-infection also diminished transplacental transfer of antibodies. Importantly, IgG1 was the most affected by maternal HIV infection but, depending on the pathogen, other subclasses were also affected. *P. falciparum* exposure also reduced the levels and transfer of some antibodies, although overall the effect was lower than HIV infection. Our findings are important for effective maternal immunization strategies and for newborn and infant’s health.

## Materials and methods

### Study design and sample collection

A total of 197 HIV-uninfected and 144 HIV-infected women were recruited among those participating in two clinical trials of antimalarial intermittent preventive treatment in pregnancy (IPTp, ClinicalTrialGov NCT00811421) (Fig. 7) in the Manhiça District, Southern Mozambique [73, 74], between May 2011 and September 2012, to perform an immunology ancillary study. The first clinical trial evaluated mefloquine (MQ) as an alternative IPTp drug to sulfadoxine-pyrimethamine (SP) in HIV-uninfected pregnant women. The study arms were (1) SP, (2) single dose MQ (MQ full), and (3) split dose over two days MQ (MQ split). The second trial evaluated MQ as IPTp drug in HIV-infected pregnant women in whom SP is contraindicated and who received daily cotrimoxazole (CTX), and women received either three monthly doses of MQ or placebo. All women received bed nets treated with long-lasting insecticide and supplements of folic acid and ferrous sulphate. Women also received tetanus vaccination. At the time of the study, the intensity of malaria transmission was low/moderate [75]. Antiretroviral therapy (ART) with daily monotherapy with zidovudine (AZT) was recommended when CD4^+^ T cell count was below <350 cells/μL and/or when women were in III or IV HIV WHO clinical stage [76].

**Fig. 7:**
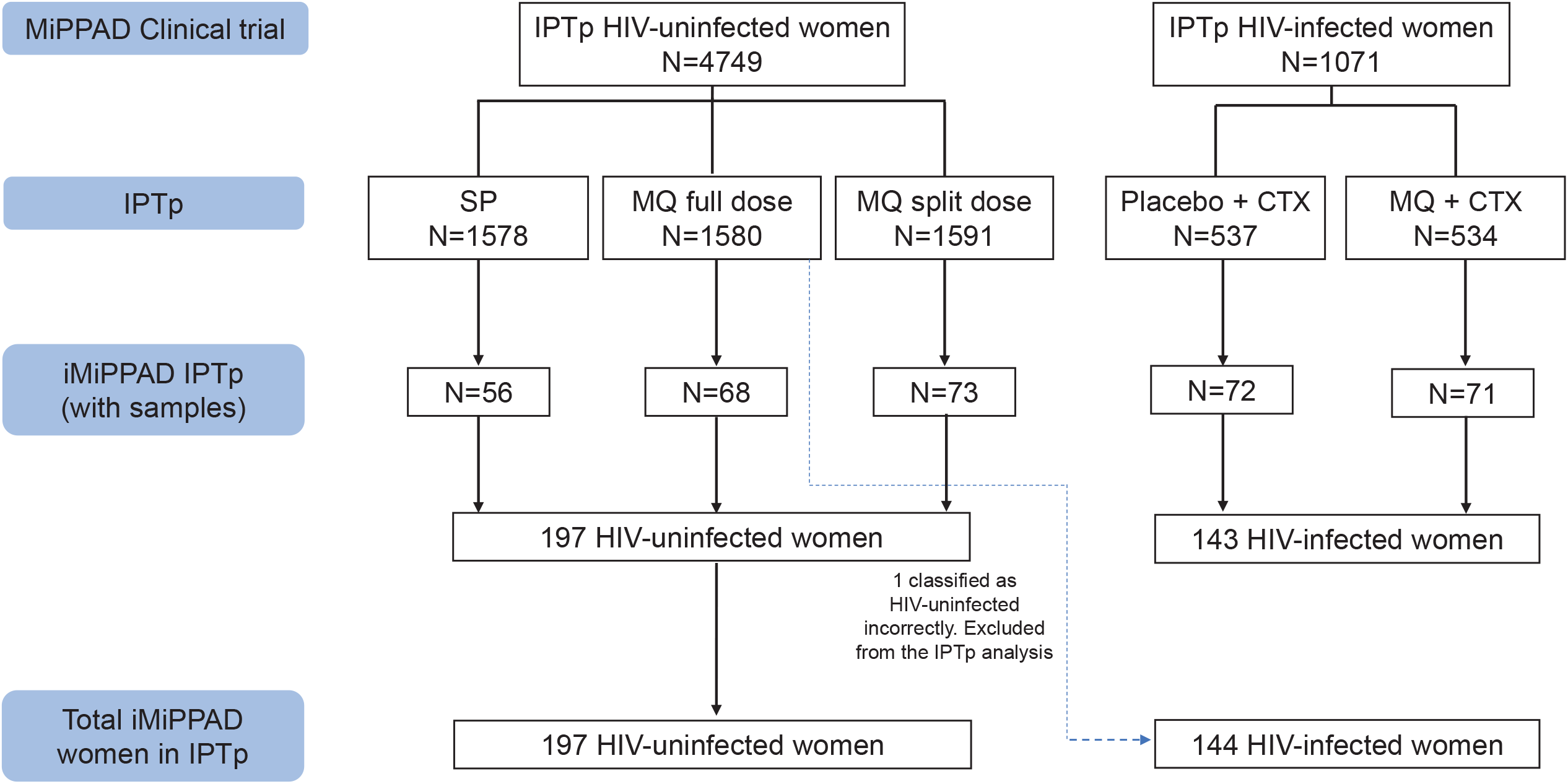
IPTp trial profile.

At delivery, blood samples from women (peripheral, placental and cord blood) were collected into sodium heparin and EDTA vacutainers. Plasma samples from peripheral blood and cord blood were available for this study from 332 (195 HIV-uninfected and 137 HIV-infected) and 303 women (178 HIV-uninfected and 125 HIV-infected), respectively. There were 294 mother-cord paired samples.

For the detection of *P. falciparum* species, thick and thin blood smears were assessed according to standard procedures [73, 74]. Fifty μl of maternal peripheral, placental, and cord blood samples were collected on filter papers for the detection of *P. falciparum* by means of a real-time quantitative polymerase-chain-reaction (qPCR) assay targeting the 18S ribosomal RNA [77]. Tissue samples from the maternal side of the placenta were also collected for the assessment of placental malaria. Microscopy data of peripheral and placental blood smears at delivery were available for 308 and 340 women, respectively. Peripheral and placental blood qPCR data were available for 242 and 236 women, respectively.

### Antibody assays

Quantitative suspension array technology (qSAT) assays applying the xMAP™ technology (Luminex Corp., TX) were used to measure antigen-specific IgG, IgG1, IgG2, IgG3 and IgG4 responses to vaccine and pathogen antigens. A total of 16 recombinant proteins were selected for the analysis: diphtheria toxoid (*Corynebacterium diphtheriae,* Alpha Diagnostic DTOX15-N-500), tetanus toxin (*Clostridium tetani,* Santa Cruz SC222347), pertussis toxin (*Bordetella pertussis,* Santa Cruz SC200837), *Hib* Oligosaccharide (BEI Resources NR12268), pneumococcal surface protein A (PspA, *Streptococcus pneumoniae,* BEI Resources NR33179), shiga toxin (*Shigella dysenteriae,* BEI Resources NR4676), anti-O-specific polysaccharide (OSP, *Vibrio cholerae,* Massachusetts General Hospital, MA, USA) [78], hepatitis B surface antigen (HBsAg, Abcam ab91276), hemagglutinin (measles, Alpha Diagnostic RP655), viral protein 6 (VP6, rotavirus, Friedzgerald 80-1389), F protein (respiratory syncytial virus, BEI Resources NR31097), 17-kDA surface antigen (Cp17, *Cryptosporidium parvum,* Centres for Disease Control and Prevention, GA, USA) [79], variant-specific surface protein 5 (VSP5, *Giardia intestinalis*) [79], 42 kDA fragment of merozoite surface protein 1 (MSP1_42_, *P. falciparum*, WRAIR) [80], merozoite surface protein 2 (MSP2, *P. falciparum,* University of Edinburgh) [81] and exported protein 1 (EXP1, *P. falciparum,* Sanaria) [82]. MSP1_42_ antigen was selected for representing *P. falciparum* infection. Eight recombinant proteins represent the most prevalent pathogens circulating in the study area [83–85] and 6 were from the vaccines administrated to the infants through the EPI in Mozambique [86].

qSAT assays were previously standardized and optimized to control for sources of variability [87–89]. Briefly, antigens covalently coupled to MagPlex beads were added to a 96-well μClear® flat bottom plate (Greiner Bio-One) in multiplex resuspended in 50μL of PBS, 1% BSA, 0.05% Azide pH 7.4 (PBS-BN). Fifty μL of test samples, negative or positive controls [90] were added to multiplex wells and incubated overnight at 4°C protected from light. After incubation, plates were washed three times with PBS-Tween 20 0.05%, and 100μL of anti-human IgG (Sigma B1140), anti-human IgG1 (Abcam ab99775), anti-human IgG2 (Invitrogen MA1-34755), anti-human IgG3 (Sigma B3523) or anti-human IgG4 (Invitrogen MA5-16716), each at their corresponding dilution, were added and incubated for 45 min. Then, plates were washed three times more and 100μL of streptavidin-R-phycoerythrin (Sigma 42250) at the appropriate dilution were added to all wells and incubated 30 min for IgG, IgG1 and IgG3. For IgG2 and IgG4, 100 μL of anti-mouse IgG (Fc-specific)−biotin (Merck B7401) were added and incubated for 45 min, followed by another washing cycle and the incubation with streptavidin-R-phycoerythrin for 30 min. Finally, plates were washed and beads resuspended in 100 μL/well of PBS-BN. Plates were read using the Luminex 100/200 analyser, and at least 20 microspheres per analyte were acquired per sample. Antibody levels were measured as median fluorescence intensity (MFI). Data were captured using xPonent software.

Test samples were assayed at 2 dilutions for IgG (1/250 and 1/10000), and IgG1 and IgG3 (1/100 and 1/2500) to ensure that at least one dilution fell in the linear range of the respective standard curve. For IgG2 and IgG4 only 1 dilution was tested (1/50) because their usual low levels. Twelve serial dilutions (1:3, starting at 1/25) of a positive control (WHO Reference Reagent for anti-malaria *P. falciparum* human serum, NIBSC code: 10/198) were used for QA/QC and to select optimal sample dilution for data analysis. Two blanks were added to each plate also for quality control purposes. Sample distribution across plates was designed to ensure a balanced distribution of groups and time-points. Single replicates of the assay were performed.

### Statistical Analysis

To stabilize the variance, the analysis was done on log_10_-transformed values of the MFI measurements. To select the sample dilution for each antigen-isotype/subclass-plate, the dilution nearest to the midpoint between the two standard curve serial dilutions ranging the maximum slope was chosen. If the maximum MFI value of a standard curve did not reach 15000, the reference value was automatically set up at 15000, since below this point, standard curve data does not seem trustworthy. If the MFI of the first sample dilution was lower than the MFI of the second dilution (hook effect), the second one was chosen. Plates were normalized using the standard curve in each plate and the average standard curve from all plates -in both cases using the dilution of the latter with the value closest to 15000 MFI. The MFI values of samples were multiplied by the corresponding normalization factor (MFI value of the chosen dilution from the average standard curve divided by the MFI value of same dilution in the plate curve).

The Shapiro-Wilk test of normality confirmed that most of the antibody data were not normally distributed. The Chi-square and the non–parametric Wilcoxon-Mann-Whitney tests were used to compare categorical and continuous variables, respectively, between HIV-infected and HIV-uninfected women. Comparisons of crude Ig levels across antigens and subclasses between HIV exposure groups were assessed by Wilcoxon-Mann-Whitney tests. Univariable linear regression models were fit to determine the effect of variables on the cord blood antibody levels (log_10_) or the cord blood/mother ratio (log_10_). The variables considered in this analysis were log10 maternal antibody levels, maternal HIV infection, *P. falciparum* exposure, PM (acute, defined by the presence of parasites on sections without malaria pigment; chronic, by presence of parasites and pigment; or past, by the presence of pigment alone), age, gravidity (defined as *primigravidae* and *multigravidae*), maternal anaemia (defined as haemoglobin level <11g/dL), low birth weight (defined as <2500g at birth), prematurity (defined as delivery before 37 weeks of gestational age), gestational age (measured by Ballard score [91]), treatment (defined as MQ or placebo in the HIV-infected women ancillary study and MQ full, MQ split or SP in HIV-uninfected women ancillary study), antiretroviral therapy (ART) received before the initiation of the study, CD4^+^ T cell counts (<350 cells/μL or ≥350 cells/μL), HIV viral load (<400, 400-999, 1000-9999 and >9999 copies/mL), and seasonality (dry or rainy). Exposure to *P. falciparum* was computed as the sum of the maternal IgG antibody levels (MFI) for the following immunogenic *P. falciparum* antigens: MSP1_42_, MSP2 and EXP1, as antibody levels to these antigens have been shown to reflect exposure to malaria [92, 93]. Seasonality was computed for each woman based on the pregnancy period - if at least 4 of the pregnancy months felt under the category of rainy period (November through April), the season was defined as such. In any other case, the season was defined as dry. A base multivariable model including maternal antibody levels, maternal HIV infection and *P. falciparum* exposure was established for each antigen and IgG or IgG subclass. Base model for MSP_42_ did not include *P. falciparum* exposure as this variable includes antibodies to this antigen. We performed additional regression models testing exhaustively all possible combinations of predictor variables (added to our base model) and selected the models based on the Akaike information criterion (AIC), Bayesian information criterion (BIC) and Adjusted r-square parameters. All p-values were considered statistically significant when <0.05 after adjusting for multiple testing through Benjamini-Hochberg. All data collected were pre-processed, managed and analysed using the R software version 3.6.3 and its package *devtools* [94]. The *ggplot2* package was used to perform boxplot graphs [95]. The *FactoMineR* and *factoextra* packages were used to perform Principal Component Analysis (PCA) [96, 97].

## Additional files

Supplementary material 1: Cord blood levels and placental transfer of antibodies univariable models.

## Acknowledgments

We are grateful to the volunteers and their families; the clinical, field, and lab teams at the from the Manhiça Health Research Centre, particularly the lab personnel Bendita Zavale, Lázaro Quimice, Elias Matusse, Eugenio Mussa and Edmundo José. Special thanks to Laura Puyol from ISGlobal for her logistic support. We thank Luis Izquierdo for the assessment in the protein expression protocol. We are grateful to Jeffrey Priest from CDC for the Cp17 and VSP5 plasmid proteins, Edward Ryan from Massachusetts General Hospital for the OSP antigen, and David Cavanagh from University of Edinburgh for the MSP2.

## Figures legends and tables

**Fig. 2-figure supplement 1:**
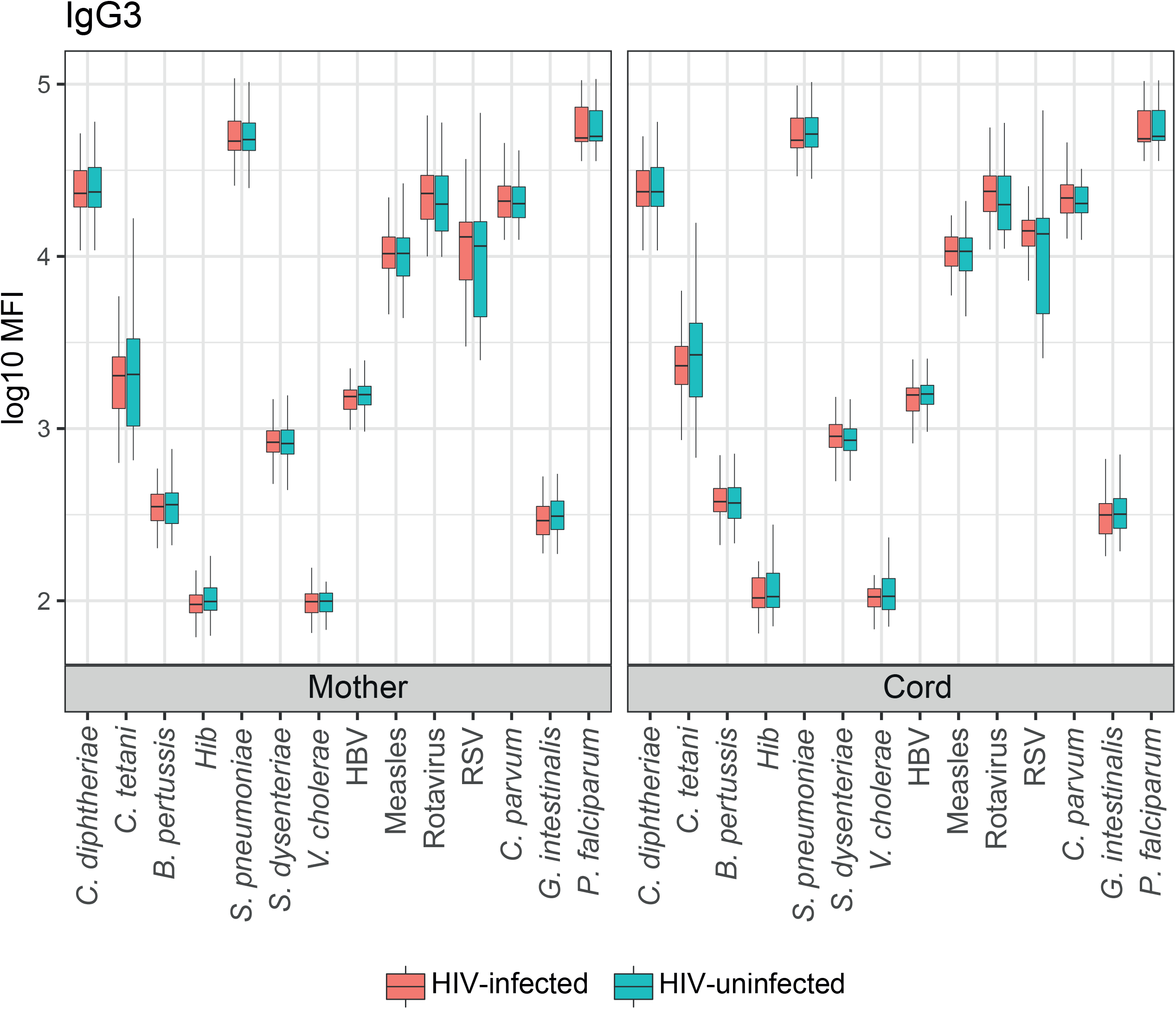
Mother and cord blood antibody levels (log_10_ MFI) in HIV-infected and HIV-uninfected women. Boxplots illustrate the medians and the interquartile range for IgG3. Levels between HIV-infected and HIV-uninfected women were compared by parametric Wilcoxon-Mann–Whitney test and p-values were adjusted for multiple testing by the Benjamini-Hochberg approach. Statistically significant differences between HIV infected and uninfected women levels are highlighted with an asterisk. Red represents HIV-infected women and blue HIV-uninfected women.

**Fig. 3-figure supplement 1:**
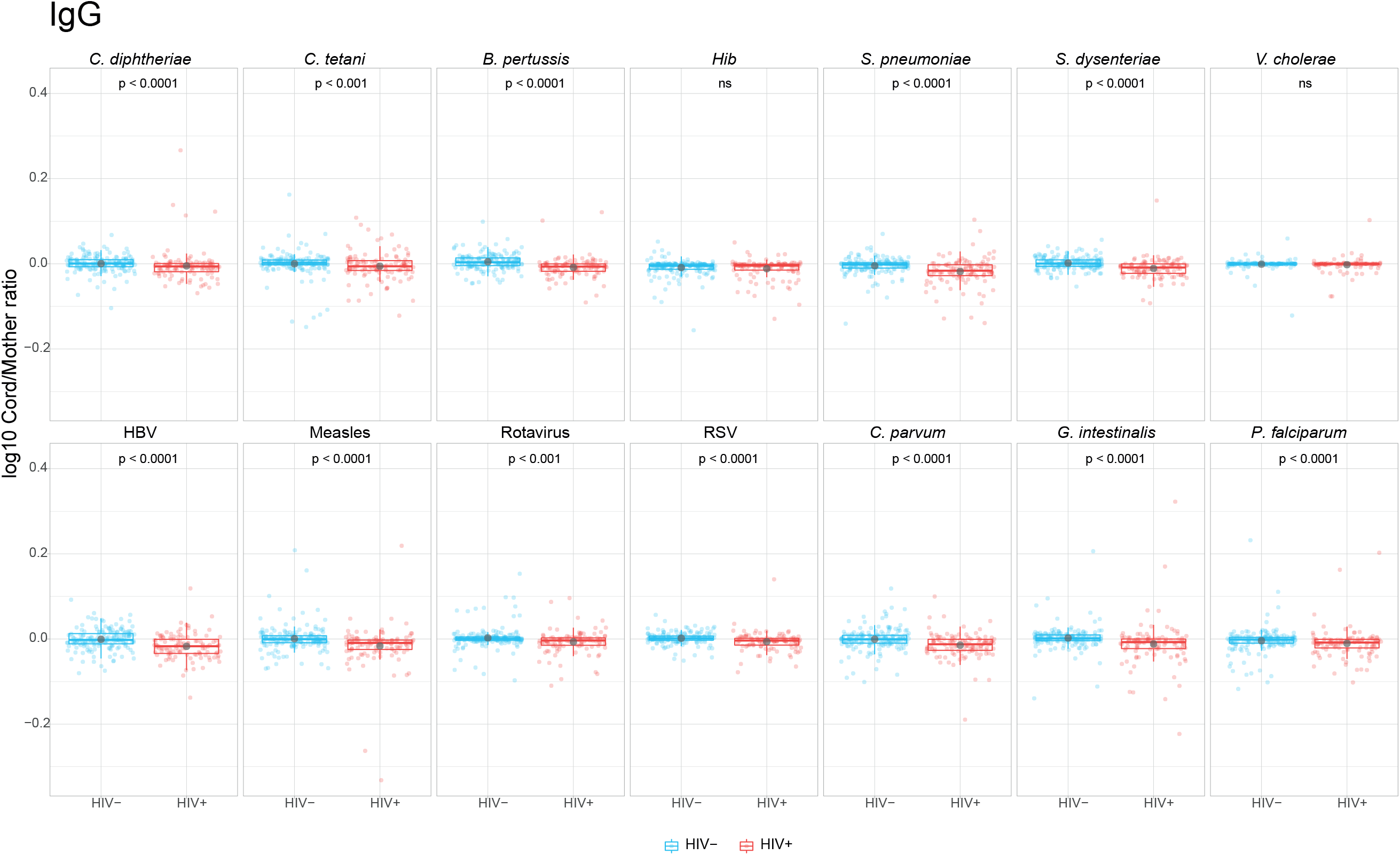
Cord/mother log_10_ antibody ratios in HIV-infected and HIV-uninfected women. Boxplots illustrate the medians, the interquartile range (IQR) and the outlier points that are further 1.5*IQR and black dots show the arithmetic means for IgG. Levels between HIV-infected and uninfected women were compared by Wilcoxon test and p-values were adjusted for multiple testing by the Benjamini-Hochberg approach. ns = not significant. Red represents HIV-infected women and blue HIV-uninfected women.

**Fig. 3-figure supplement 2:**
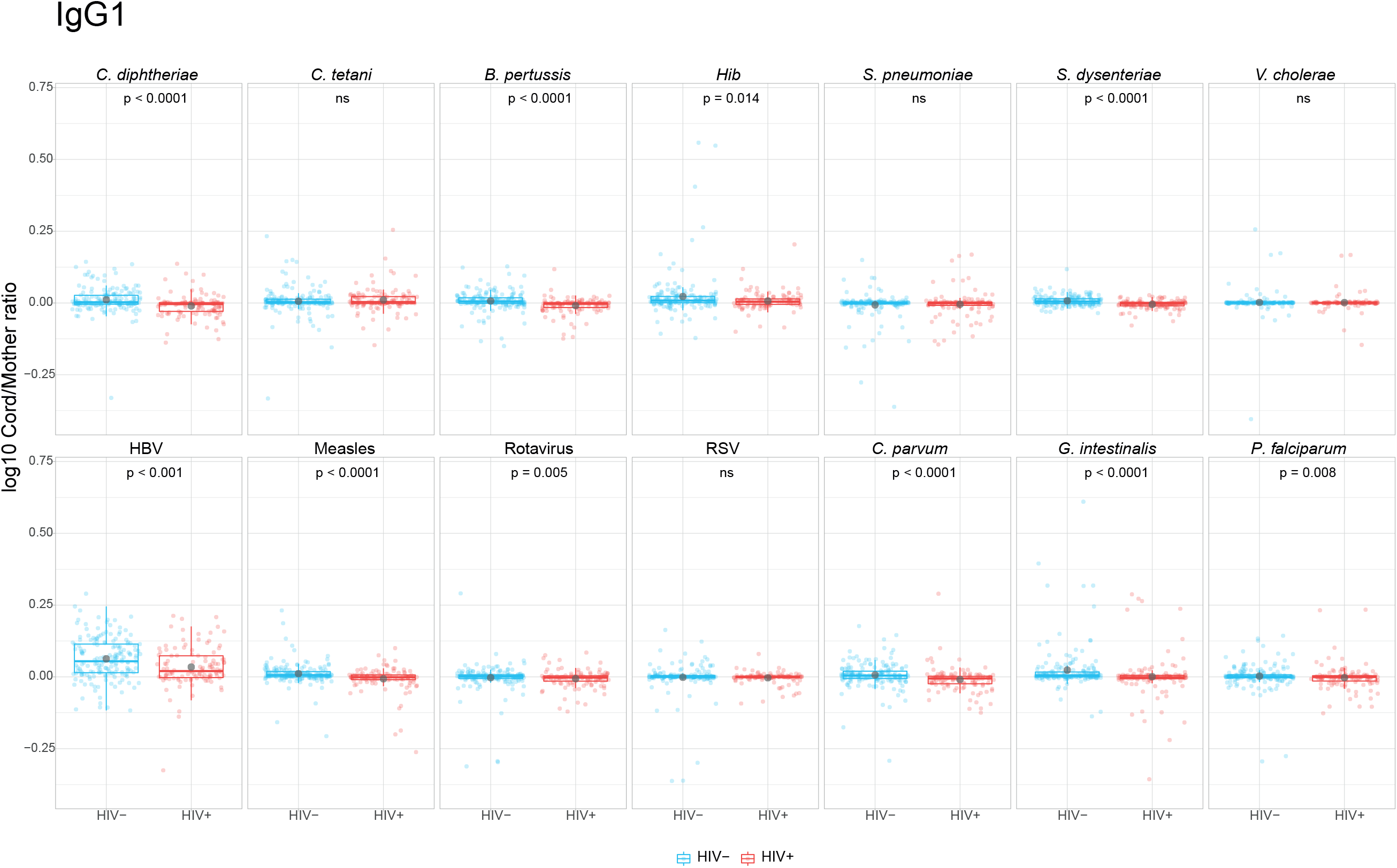
Cord/mother log_10_ antibody ratios in HIV-infected and HIV-uninfected women. Boxplots illustrate the medians, the interquartile range (IQR) and the outlier points that are further 1.5*IQR and black dots show the arithmetic means for IgG1. Levels between HIV-infected and uninfected women were compared by Wilcoxon test and p-values were adjusted for multiple testing by the Benjamini-Hochberg approach. ns = not significant. Red represents HIV-infected women and blue HIV-uninfected women.

**Fig. 3-figure supplement 3:**
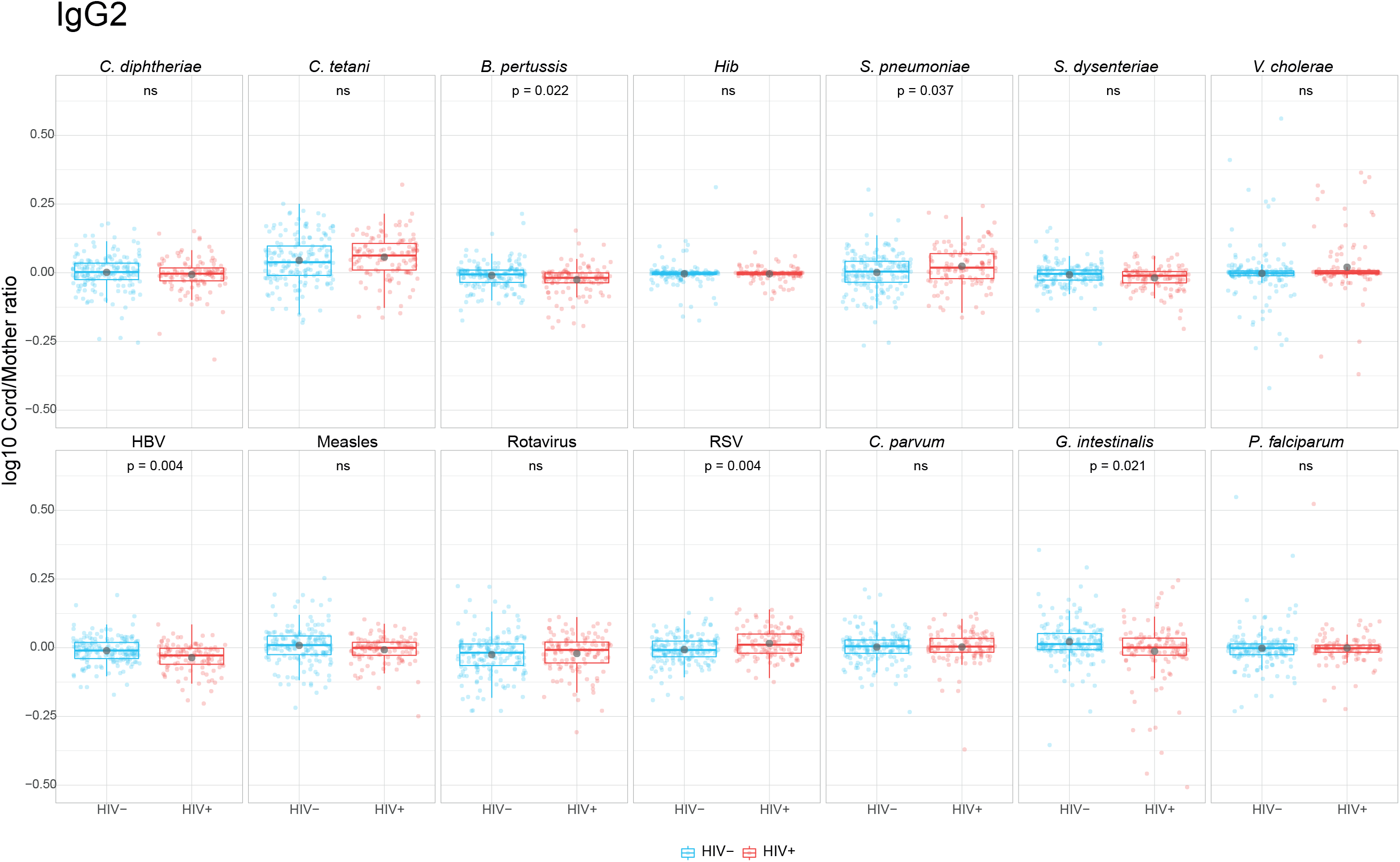
Cord/mother log_10_ antibody ratios in HIV-infected and HIV-uninfected women. Boxplots illustrate the medians, the interquartile range (IQR) and the outlier points that are further 1.5*IQR and black dots show the arithmetic means for IgG2. Levels between HIV-infected and uninfected women were compared by Wilcoxon test and p-values were adjusted for multiple testing by the Benjamini-Hochberg approach. ns = not significant. Red represents HIV-infected women and blue HIV-uninfected women.

**Fig. 3-figure supplement 4:**
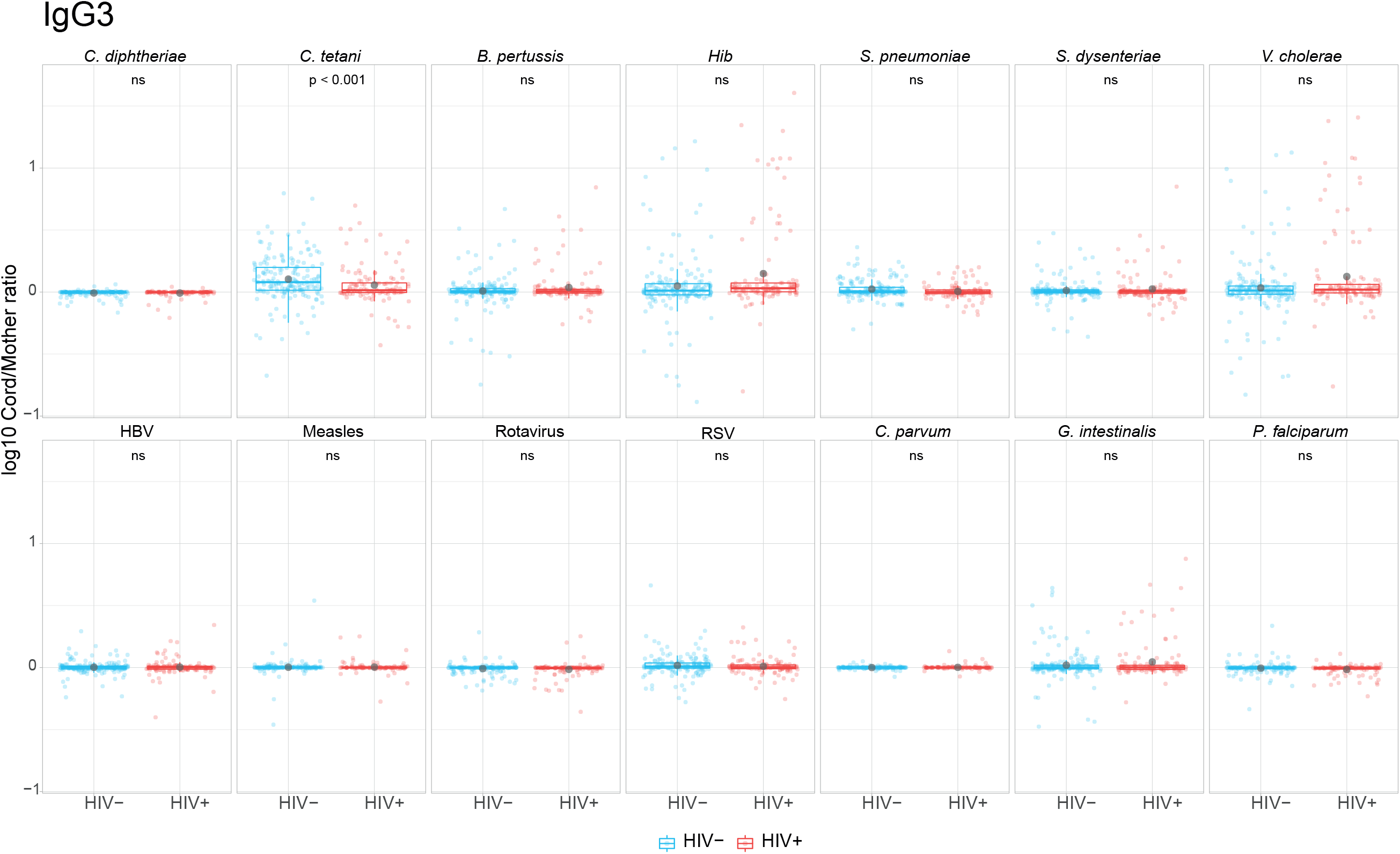
Cord/mother log_10_ antibody ratios in HIV-infected and HIV-uninfected women. Boxplots illustrate the medians, the interquartile range (IQR) and the outlier points that are further 1.5*IQR and black dots show the arithmetic means for IgG3. Levels between HIV-infected and uninfected women were compared by Wilcoxon test and p-values were adjusted for multiple testing by the Benjamini-Hochberg approach. ns = not significant. Red represents HIV-infected women and blue HIV-uninfected women.

**Fig.3-figure supplement 5:**
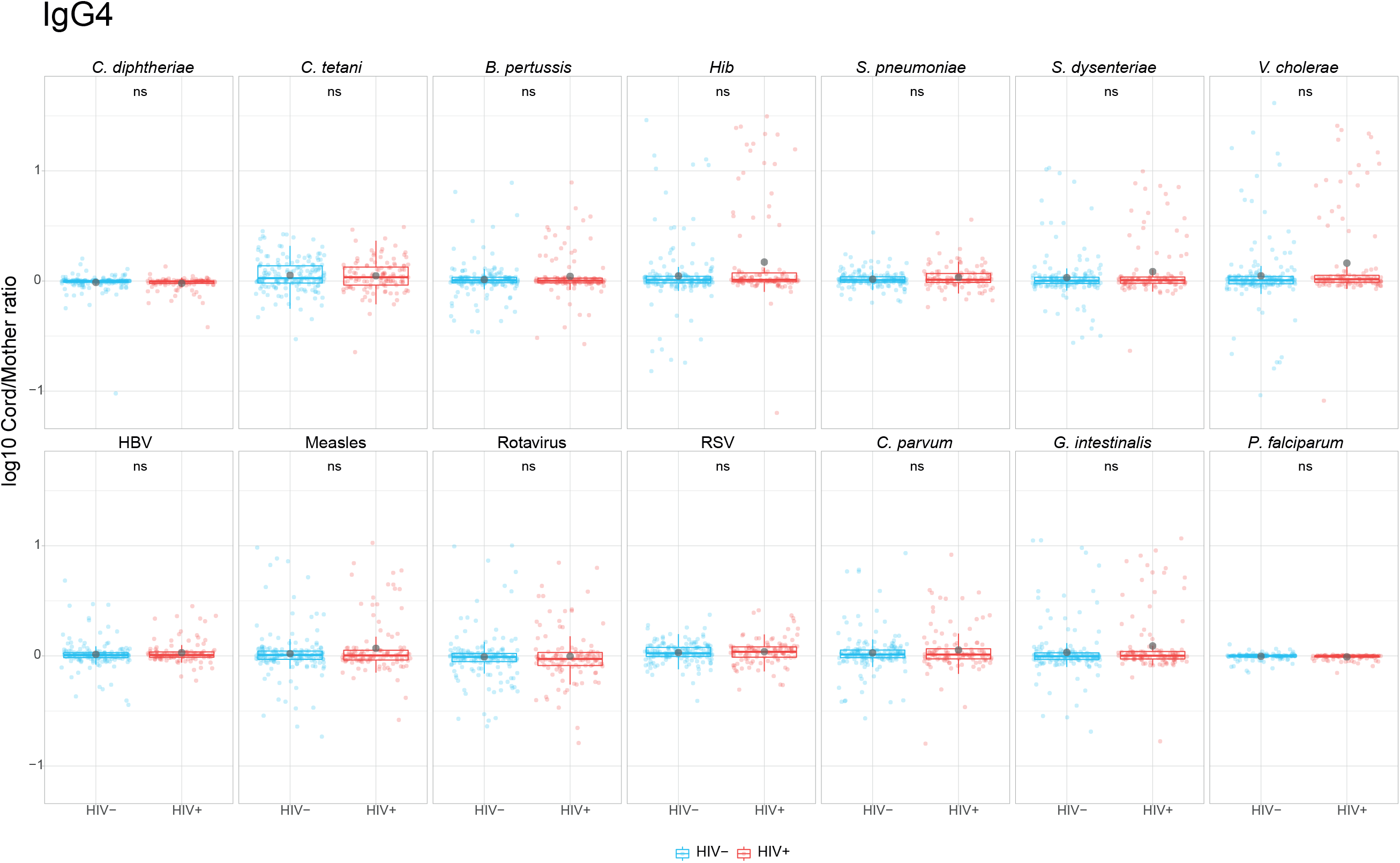
Cord/mother log_10_ antibody ratios in HIV-infected and HIV-uninfected women. Boxplots illustrate the medians, the interquartile range (IQR) and the outlier points that are further 1.5*IQR and black dots show the arithmetic means for IgG4. Levels between HIV-infected and uninfected women were compared by Wilcoxon test and p-values were adjusted for multiple testing by the Benjamini-Hochberg approach. ns = not significant. Red represents HIV-infected women and blue HIV-uninfected women.

**Fig.4-figure supplement 1:**
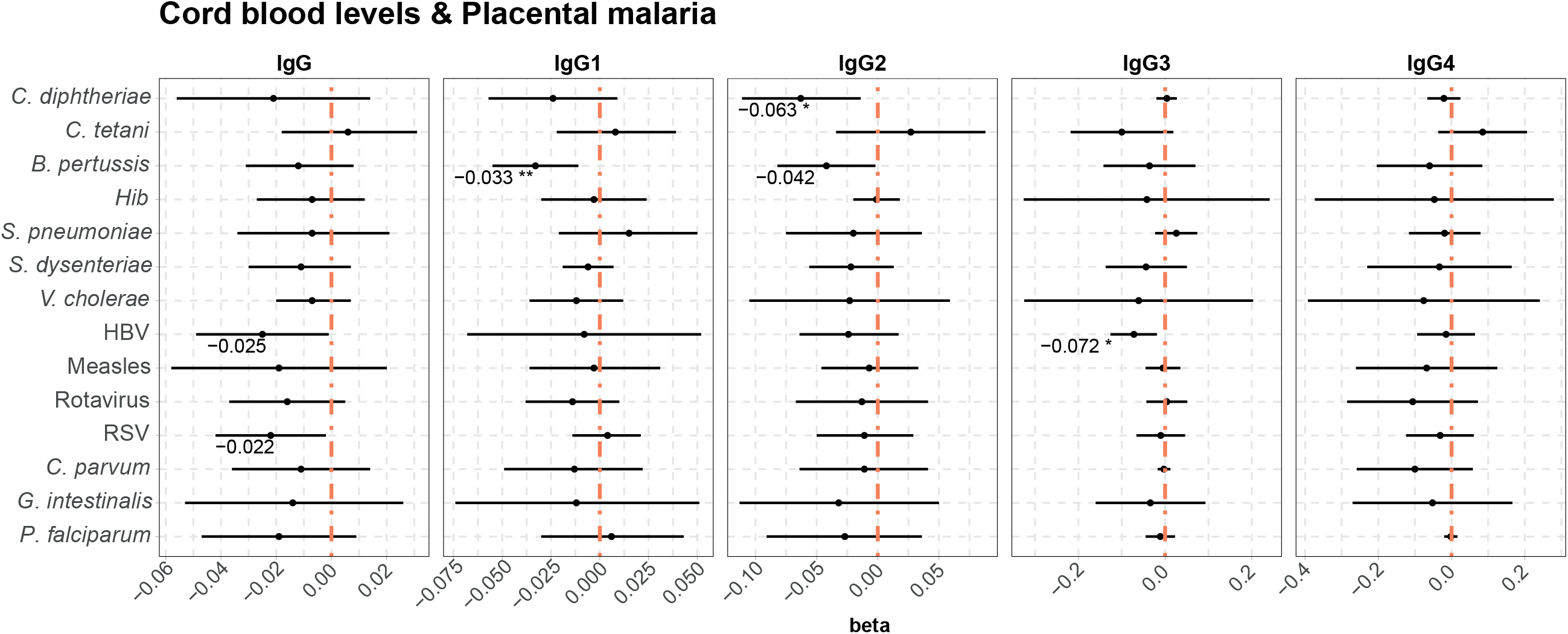
Forest plots show the effect of placental malaria on cord blood levels of IgG and IgG subclasses, for all the antigens tested, in HIV-infected women. Cord antibody levels are represented in log_10_MFI. Beta values are shown when raw p-vals are significant. Asterisks are shown when adjusted p-vals by Benjamini-Hochbert are significant. **** = p-val < 0.0001, *** = p-val < 0.001, ** = p-val < 0.01, * = p-val < 0.05.

**Fig. 6-figure supplement 1:**
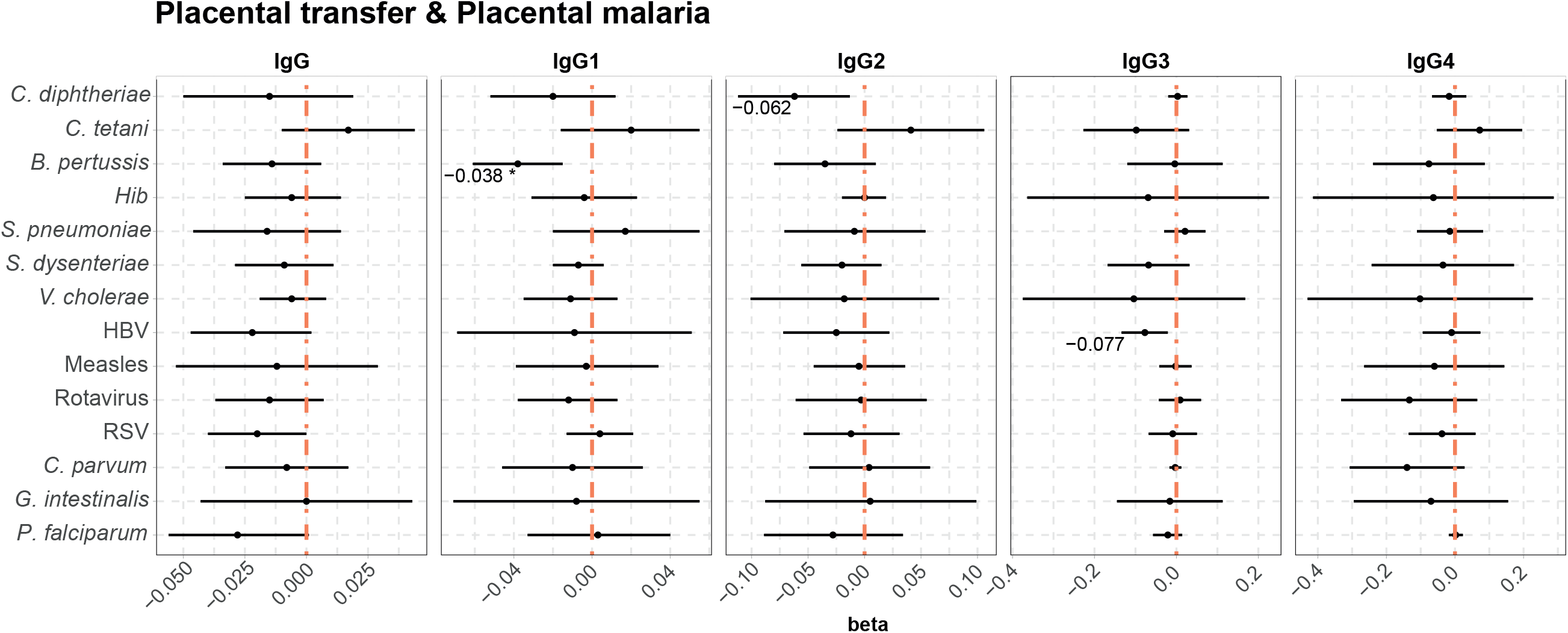
Forest plots show the effect of placental malaria on placental transfer of IgG and IgG subclasses, for all the antigens tested, in HIV-infected women. Placental transfer is represented as cord/mother ratio (log_10_). Beta values are shown when raw p-vals are significant. Asterisks are shown when adjusted p-vals by Benjamini-Hochbert are significant. **** = p-val < 0.0001, *** = p-val < 0.001, ** = p-val < 0.01, * = p-val < 0.05.

## References

[1] World Health Organization. World Health Statistics 2018: Monitoring health for the SDGs. 2018.

[2] WHO | Infant mortality. WHO, https://www.who.int/gho/child_health/mortality/neonatal_infant_text/en/ (2018, accessed 27 November 2019).

[3] Kollmann TR, Kampmann B, Mazmanian SK, et al. Protecting the Newborn and Young Infant from Infectious Diseases: Lessons from Immune Ontogeny. Immunity 2017; 46: 350–363.

[4] Basha S, Surendran N, Pichichero M. Immune responses in neonates. Expert Review of Clinical Immunology 2014; 10: 1171–1184.

[5] Prabhudas M, Adkins B, Gans H, et al. Challenges in infant immunity: Implications for responses to infection and vaccines. Nature Immunology 2011; 12: 189–194.

[6] Zhang X, Zhivaki D, Lo-Man R. Unique aspects of the perinatal immune system. Nat Rev Immunol 2017; 17: 495–507.

[7] O’Brien KL, Baggett HC, Brooks WA, et al. Causes of severe pneumonia requiring hospital admission in children without HIV infection from Africa and Asia: the PERCH multi-country case-control study. Lancet 2019; 394: 757–779.

[8] Kabego L, Balol’Ebwami S, Kasengi JB, et al. Human respiratory syncytial virus: Prevalence, viral co-infections and risk factors for lower respiratory tract infections in children under 5 years of age at a general hospital in the democratic republic of Congo. J Med Microbiol 2018; 67: 514–522.

[9] Palmeira P, Quinello C, Silveira-Lessa AL, et al. IgG placental transfer in healthy and pathological pregnancies. Clinical and Developmental Immunology; 2012. Epub ahead of print 2012. DOI: 10.1155/2012/985646.

[10] Wood N, Siegrist CA. Neonatal immunization: Where do we stand? Curr Opin Infect Dis 2011; 24: 190–195.

[11] Morris MC, Surendran N. Neonatal vaccination: Challenges and intervention strategies. Neonatology 2016; 109: 161–169.

[12] Demirjian A, Levy O. Safety and efficacy of neonatal vaccination. European Journal of Immunology 2009; 39: 36–46.

[13] Deogaonkar R, Hutubessy R, van der Putten I, et al. Systematic review of studies evaluating the broader economic impact of vaccination in low and middle income countries. BMC Public Health 2012; 12: 878.

[14] Clemens J, Holmgren J, Kaufmann SHE, et al. Ten years of the global Alliance for vaccines and immunization: Challenges and progress. Nature Immunology 2010; 11: 1069–1072.

[15] Saso A, Kampmann B. Vaccine responses in newborns. Seminars in Immunopathology 2017; 39: 627–642.

[16] Simister NE, Story CM, Chen H-L, et al. An IgG-transporting Fc receptor expressed in the syncytiotrophoblast of human placenta. Eur J Immunol 1996; 26: 1527–1531.

[17] Malek A, Sager R, Kuhn P, et al. Evolution of Maternofetal Transport of Immunoglobulins During Human Pregnancy. Am J Reprod Immunol 1996; 36: 248–255.

[18] Bundhoo A, Paveglio S, Rafti E, et al. Evidence that FcRn mediates the transplacental passage of maternal IgE in the form of IgG anti-IgE/IgE immune complexes. Clin Exp Allergy 2015; 45: 1085–1098.

[19] Ferrante A, Beard LJ, Feldman RG. IgG subclass distribution of antibodies to bacteria and viral antigens. Pediatr Infect Dis J 1990; 9: S16–S24.

[20] Costa-Carvalho BT, Vieira HM, Dimantas RBR, et al. Transfer of IgG subclasses across placenta in term and preterm newborns. Brazilian J Med Biol Res 1996; 29: 201–204.

[21] Marchant A, Sadarangani M, Garand M, et al. Maternal immunisation: collaborating with mother nature. The Lancet Infectious Diseases 2017; 17: e197–e208.

[22] Buchy P, Badur S, Kassianos G, et al. Vaccinating pregnant women against influenza needs to be a priority for all countries: An expert commentary. International Journal of Infectious Diseases 2020; 92: 1–12.

[23] Thwaites CL, Beeching NJ, Newton CR. Maternal and neonatal tetanus. In: The Lancet. Lancet Publishing Group, 2015, pp. 362–370.

[24] Gkentzi D, Katsakiori P, Marangos M, et al. Maternal vaccination against pertussis: A systematic review of the recent literature. Archives of Disease in Childhood: Fetal and Neonatal Edition 2017; 102: F456–F463.

[25] Heath PT, Culley FJ, Jones CE, et al. Group B streptococcus and respiratory syncytial virus immunisation during pregnancy: a landscape analysis. The Lancet Infectious Diseases 2017; 17: e223–e234.

[26] Cumberland P, Shulman CE, Maple PA, et al. Maternal HIV infection and placental malaria reduce transplacental antibody transfer and tetanus antibody levels in newborns in Kenya. J Infect Dis 2007; 196: 550–557.

[27] Jones CE, Kampmann B, Hesseling A. Maternal HIV infection and antibody responses in uninfected infants: In reply. JAMA - Journal of the American Medical Association 2011; 305: 1964–1965.

[28] Caceres VM, Strebel PM, Sutter RW. Factors Determining Prevalence of Maternal Antibody to Measles Virus throughout Infancy: A Review. Clin Infect Dis 2000; 31: 110–119.

[29] Wilcox CR, Holder B, Jones CE. Factors affecting the FcRn-mediated transplacental transfer of antibodies and implications for vaccination in pregnancy. Frontiers in Immunology; 8. Epub ahead of print 13 October 2017. DOI: 10.3389/fimmu.2017.01294.

[30] Van Den Berg JP, Westerbeek EAM, Berbers GAM, et al. Transplacental transport of IgG antibodies specific for pertussis, diphtheria, tetanus, haemophilus influenzae type b, and neisseria meningitidis serogroup C is lower in preterm compared with term infants. Pediatr Infect Dis J 2010; 29: 801–805.

[31] De Moraes-Pinto MI, Verhoeff F, Chimsuku L, et al. Placental antibody transfer: Influence of maternal HIV infection and placental malaria. Arch Dis Child Fetal Neonatal Ed; 79. Epub ahead of print 1998. DOI: 10.1136/fn.79.3.F202.

[32] Brair M-E, Brabin B, Milligan P, et al. Reduced transfer of tetanus antibodies with placental malaria. Lancet 1994; 343: 208–209.

[33] Okoko BJ, Wesuperuma LH, Ota MO, et al. Influence of placental malaria infection and maternal hypergammaglobulinaemia on materno-foetal transfer of measles and tetanus antibodies in a rural west African population. J Health Popul Nutr 2001; 19: 59–65.

[34] Atwell JE, Thumar B, Robinson LJ, et al. Impact of placental malaria and hypergammaglobulinemia on transplacental transfer of respiratory syncytial virus antibody in Papua New Guinea. J Infect Dis 2016; 213: 423–431.

[35] Okoko BJ, Wesumperuma LH, Ota MO, et al. The influence of placental malaria infection and maternal hypergammaglobulinemia on transplacental transfer of antibodies and IgG subclasses in a rural West African population. J Infect Dis 2001; 184: 627–632.

[36] Ray JE, Dobbs KR, Ogolla SO, et al. Reduced Transplacental Transfer of Antimalarial Antibodies in Kenyan HIV-Exposed Uninfected Infants. Open Forum Infect Dis; 6. Epub ahead of print 1 June 2019. DOI: 10.1093/ofid/ofz237.

[37] Scott S, Cumberland P, Shulman CE, et al. Neonatal Measles Immunity in Rural Kenya: The Influence of HIV and Placental Malaria Infections on Placental Transfer of Antibodies and Levels of Antibody in Maternal and Cord Serum Samples. J Infect Dis 2005; 191: 1854–1860.

[38] de Moraes-Pinto MI, Almeida AC, Kenj G, et al. Placental transfer and maternally acquired neonatal IgG immunity in human immunodeficiency virus infection. J Infect Dis 1996; 173: 1077–84.

[39] Moro L, Bardaji A, Nhampossa T, et al. Malaria and HIV Infection in Mozambican Pregnant Women Are Associated With Reduced Transfer of Antimalarial Antibodies to Their Newborns. J Infect Dis 2015; 211: 1004–1014.

[40] Jones CE, Naidoo S, De Beer C, et al. Maternal HIV infection and antibody responses against vaccine-preventable diseases in uninfected infants. JAMA - J Am Med Assoc 2011; 305: 576–584.

[41] Jones C, Pollock L, Barnett SM, et al. Specific antibodies against vaccine-preventable infections: A mother-infant cohort study. BMJ Open; 3. Epub ahead of print 2013. DOI: 10.1136/bmjopen-2012-002473.

[42] Dechavanne C, Cottrell G, Garcia A, et al. Placental Malaria: Decreased transfer of maternal antibodies directed to Plasmodium falciparum and impact on the incidence of febrile infections in infants. PLoS One; 10. Epub ahead of print 1 December 2015. DOI: 10.1371/journal.pone.0145464.

[43] Boudová S, Divala T, Mungwira R, et al. Placental but Not Peripheral Plasmodium falciparum Infection During Pregnancy Is Associated With Increased Risk of Malaria in Infancy. In: Journal of Infectious Diseases. 2017. Epub ahead of print 2017. DOI: 10.1093/infdis/jix372.

[44] van den Berg JP, Westerbeek EAM, van der Klis FRM, et al. Transplacental transport of IgG antibodies to preterm infants: A review of the literature. Early Human Development 2011; 87: 67–72.

[45] Fu C, Lu L, Wu H, et al. Placental antibody transfer efficiency and maternal levels: Specific for measles, coxsackievirus A16, enterovirus 71, poliomyelitis I-III and HIV-1 antibodies. Sci Rep 2016; 6: 1–6.

[46] Pou C, Nkulikiyimfura D, Henckel E, et al. The repertoire of maternal anti-viral antibodies in human newborns. Nat Med 2019; 25: 591–596.

[47] Askari A, Hakimi H, Nasiri Ahmadabadi B, et al. Prevalence of hepatitis B co-infection among HIV positive patients: Narrative review article. Iranian Journal of Public Health 2014; 43: 705–712.

[48] Feitosa G, Bandeira AC, Sampaio DP, et al. High prevalence of giardiasis and stronglyloidiasis among HIV-infected patients in Bahia, Brazil. Braz J Infect Dis 2001; 5: 339–344.

[49] Angarano G, Maggi P, Di Bari MA, et al. Giardiasis in hiv: A possible role in patients with severe immune deficiency. Eur J Epidemiol 1997; 13: 485–487.

[50] De Milito A, Nilsson A, Titanji K, et al. Mechanisms of hypergammaglobulinemia and impaired antigen-specific humoral immunity in HIV-1 infection. Blood 2004; 103: 2180–2186.

[51] Gupta A, Mathad JS, Yang WT, et al. Maternal pneumococcal capsular IgG antibodies and transplacental transfer are low in South Asian HIV-infected mother-infant pairs. Vaccine 2014; 32: 1466–1472.

[52] Jallow S, Agosti Y, Kgagudi P, et al. Impaired Transplacental Transfer of Respiratory Syncytial Virus–neutralizing Antibodies in Human Immunodeficiency Virus–infected Versus –uninfected Pregnant Women. Clin Infect Dis 2019; 69: 151–154.

[53] Weinberg A, Mussi-Pinhata MM, Yu Q, et al. Excess respiratory viral infections and low antibody responses among HIV-exposed, uninfected infants. AIDS 2017; 31: 669–679.

[54] Vidarsson G, Dekkers G, Rispens T. IgG subclasses and allotypes: From structure to effector functions. Front Immunol 2014; 5: 520.

[55] Irani V, Guy AJ, Andrew D, et al. Molecular properties of human IgG subclasses and their implications for designing therapeutic monoclonal antibodies against infectious diseases. Molecular Immunology 2015; 67: 171–182.

[56] Einhorn MS, Granoff DM, Nahm MH, et al. Concentrations of antibodies in paired maternal and infant sera: Relationship to IgG subclass. J Pediatr 1987; 111: 783–788.

[57] Seppälä I, Sarvas H, Mäkelä O, et al. Human Antibody Responses to Two Conjugate Vaccines of Haemophilus influenzae Type B Saccharides and Diphtheria Toxin. Scand J Immunol 1988; 28: 471–479.

[58] Parkkali T, Käyhty H, Anttila M, et al. IgG subclasses and avidity of antibodies to polysaccharide antigens in allogeneic BMT recipients after vaccination with pneumococcal polysaccharide and Haemophilus influenzae type b conjugate vaccines. Bone Marrow Transplant 1999; 24: 671–678.

[59] Bosire R, Farquhar C, Nduati R, et al. Higher Transplacental Pathogen-Specific Antibody Transfer among Pregnant Women Randomized to Triple Antiretroviral Treatment Versus Short Course Zidovudine. Pediatr Infect Dis J 2018; 37: 246–252.

[60] Baroncelli S, Galluzzo CM, Mancinelli S, et al. Antibodies against pneumococcal capsular polysaccharide in Malawian HIV-positive mothers and their HIV-exposed uninfected children. Infect Dis (Auckl) 2016; 48: 317–321.

[61] Farquhar C, Nduati R, Haigwood N, et al. High maternal HIV-1 viral load during pregnancy is associated with reduced placental transfer of measles IgG antibody. J Acquir Immune Defic Syndr 2005; 40: 494–497.

[62] Goetghebuer T, Smolen KK, Adler C, et al. Initiation of Antiretroviral Therapy Before Pregnancy Reduces the Risk of Infection-related Hospitalization in Human Immunodeficiency Virus-exposed Uninfected Infants Born in a High-income Country. Clin Infect Dis. Epub ahead of print 2019. DOI: 10.1093/cid/ciy673.

[63] Wesumperuma HL, Perera AJ, Pharoah POD, et al. The influence of prematurity and low birthweight on transplacental antibody transfer in Sri Lanka. Ann Trop Med Parasitol 1999; 93: 169–177.

[64] Okoko BJ, Wesumperuma LH, Hart AC. Materno-foetal transfer of H. influenzae and pneumococcal antibodies is influenced by prematurity and low birth weight: implications for conjugate vaccine trials. Vaccine 2001; 20: 647–50.

[65] Messeret ES, Masresha B, Yakubu A, et al. Maternal and Neonatal Tetanus Elimination (MNTE) in The WHO African Region. J Immunol Sci 2018; Suppl: 103–107.

[66] Zackrisson G, Lagergård T, Trollfors B. Subclass compositions of immunoglobulin G to pertussis toxin in patients with whooping cough, in healthy individuals, and in recipients of a pertussis toxoid vaccine. J Clin Microbiol 1989; 27: 1567–71.

[67] Granström M, Ferngren H, Linde A, et al. IgG subclass responses to Bordetella pertussis filamentous haemagglutinin and pertussis toxin in whooping cough. Serodiagn Immunother Infect Dis 1989; 3: 403–412.

[68] Munoz FM. Respiratory syncytial virus in infants: Is maternal vaccination a realistic strategy? Current Opinion in Infectious Diseases. Epub ahead of print 2015. DOI: 10.1097/QCO.0000000000000161.

[69] Wagner DK, Muelenaer P, Henderson FW, et al. Serum immunoglobulin G antibody subclass response to respiratory syncytial virus F and G glycoproteins after first, second, and third infections. J Clin Microbiol 1989; 27: 589–92.

[70] Higgins D, Trujillo C, Keech C. Advances in RSV vaccine research and development - A global agenda. Vaccine. Epub ahead of print 2016. DOI: 10.1016/j.vaccine.2016.03.109.

[71] Piedra PA, Jewell AM, Cron SG, et al. Correlates of immunity to respiratory syncytial virus (RSV) associated-hospitalization: Establishment of minimum protective threshold levels of serum neutralizing antibodies. In: Vaccine. Elsevier BV, 2003, pp. 3479–3482.

[72] Connor EM. Palivizumab, a humanized respiratory syncytial virus monoclonal antibody, reduces hospitalization from respiratory syncytial virus infection in high-risk infants. Pediatrics 1998; 102: 531–537.

[73] González R, Mombo-Ngoma G, Ouédraogo S, et al. Intermittent Preventive Treatment of Malaria in Pregnancy with Mefloquine in HIV-Negative Women: A Multicentre Randomized Controlled Trial. PLoS Med 2014; 11: e1001733.

[74] González R, Desai M, Macete E, et al. Intermittent Preventive Treatment of Malaria in Pregnancy with Mefloquine in HIV-Infected Women Receiving Cotrimoxazole Prophylaxis: A Multicenter Randomized Placebo-Controlled Trial. PLoS Med 2014; 11: e1001735.

[75] Mayor A, Bardají A, Macete E, et al. Changing Trends in P. falciparum Burden, Immunity, and Disease in Pregnancy. N Engl J Med 2015; 373: 1607–17.

[76] WHO. Interim WHO Clinical Staging of HIV/AIDS and HIV/AIDS Case definitions for surveillance. African Region. 2005. Epub ahead of print 2005. DOI: 10.1300/j187v04n01_05.

[77] Mayor A, Serra-Casas E, Bardají A, et al. Sub-microscopic infections and long-term recrudescence of Plasmodium falciparum in Mozambican pregnant women. Malar J 2009; 8: 1–10.

[78] Sayeed MA, Bufano MK, Xu P, et al. A Cholera Conjugate Vaccine Containing O-specific Polysaccharide (OSP) of V. cholerae O1 Inaba and Recombinant Fragment of Tetanus Toxin Heavy Chain (OSP:rTTHc) Induces Serum, Memory and Lamina Proprial Responses against OSP and Is Protective in Mice. PLoS Negl Trop Dis 2015; 9: e0003881.

[79] Priest JW, Moss DM, Visvesvara GS, et al. Multiplex assay detection of immunoglobulin G antibodies that recognize Giardia intestinalis and Cryptosporidium parvum antigens. Clin Vaccine Immunol 2010; 17: 1695–1707.

[80] Angov E, Aufiero BM, Turgeon AM, et al. Development and pre-clinical analysis of a Plasmodium falciparum Merozoite Surface Protein-142 malaria vaccine. Mol Biochem Parasitol 2003; 128: 195–204.

[81] Metzger WG, Okenu DMN, Cavanagh DR, et al. Serum IgG3 to the Plasmodium falciparum merozoite surface protein 2 is strongly associated with a reduced prospective risk of malaria. Parasite Immunol 2003; 25: 307–312.

[82] Doolan DL, Hedstrom RC, Rogers WO, et al. Identification and characterization of the protective hepatocyte erythrocyte protein 17 dDa gene of Plasmodium yoelii, homolog of Plasmodium falciparum exported protein 1. J Biol Chem 1996; 271: 17861–17868.

[83] Dgedge M, Novoa A, Macassa G, et al. The burden of disease in Maputo City, Mozambique: registered and autopsied deaths in 1994. Bull World Health Organ 2001; 79: 546–52.

[84] Kotloff KL, Nataro JP, Blackwelder WC, et al. Burden and aetiology of diarrhoeal disease in infants and young children in developing countries (the Global Enteric Multicenter Study, GEMS): A prospective, case-control study. Lancet 2013; 382: 209–222.

[85] Troeger C, Blacker B, Khalil IA, et al. Estimates of the global, regional, and national morbidity, mortality, and aetiologies of lower respiratory infections in 195 countries, 1990–2016: a systematic analysis for the Global Burden of Disease Study 2016. Lancet Infect Dis 2018; 18: 1191–1210.

[86] Lanaspa M, Balcells R, Sacoor C, et al. The performance of the expanded programme on immunization in a rural area of Mozambique. Acta Trop 2015; 149: 262–266.

[87] Vidal M, Aguilar R, Campo JJ, et al. Development of quantitative suspension array assays for six immunoglobulin isotypes and subclasses to multiple Plasmodium falciparum antigens. J Immunol Methods 2018; 455: 41–54.

[88] Ubillos I, Jiménez A, Vidal M, et al. Optimization of incubation conditions of Plasmodium falciparum antibody multiplex assays to measure IgG, IgG1-4, IgM and IgE using standard and customized reference pools for sero-epidemiological and vaccine studies. Malar J; 17. Epub ahead of print 1 June 2018. DOI: 10.1186/s12936-018-2369-3.

[89] Ubillos I, Aguilar R, Sanz H, et al. Analysis of factors affecting the variability of a quantitative suspension bead array assay measuring IgG to multiple Plasmodium antigens. PLoS One; 13. Epub ahead of print 1 July 2018. DOI: 10.1371/journal.pone.0199278.

[90] Bryan D, Silva N, Rigsby P, et al. The establishment of a WHO Reference Reagent for anti-malaria (Plasmodium falciparum) human serum. Malar J; 16. Epub ahead of print 5 August 2017. DOI: 10.1186/s12936-017-1958-x.

[91] Ballard JL, Khoury JC, Wedig K, et al. New Ballard Score, expanded to include extremely premature infants. J Pediatr 1991; 119: 417–423.

[92] Weiss GE, Traore B, Kayentao K, et al. The plasmodium falciparum-specific human memory b cell compartment expands gradually with repeated malaria infections. PLoS Pathog 2010; 6: 1–13.

[93] Dobaño C, Ubillos I, Jairoce C, et al. RTS,S/AS01E immunization increases antibody responses to vaccine-unrelated Plasmodium falciparum antigens associated with protection against clinical malaria in African children: a case-control study. BMC Med 2019; 17: 157.

[94] Wickham H, Hester J, Chang W. devtools: Tools to make developing R packages easier. R package devtools version 2.3.0., https://cran.r-project.org/package=devtools (2020, accessed 7 July 2020).

[95] Wickham H. ggplot2: Elegant Graphics for Data Analysis. Springer.

[96] Lê S, Josse J, Husson F. FactoMineR: An R package for multivariate analysis. J Stat Softw 2008; 25: 1–18.

[97] Kassambara A, Mundt F. factoextra: Extract and Visualize the Results of Multivariate Data Analyses. R package version 1.0.7., https://cran.r-project.org/package=factoextra (2020).

